# A CRISPR/Cas9-induced restoration of bioluminescence reporter system for single-cell gene expression analysis in plants

**DOI:** 10.1101/2025.05.27.656507

**Authors:** Ryohei Ueno, Shogo Ito, Tokitaka Oyama

## Abstract

Bioluminescence monitoring techniques have greatly contributed to revealing a variety of biological regulatory systems in living organisms, including circadian clocks. In plant science, these techniques are applied to long-term quantitative analyses of gene expression behavior. Transient transfection with a *luciferase* reporter using the particle bombardment method has been used for bioluminescence observations at the single-cell level. This allows for capturing heterogeneity and temporal fluctuations in cellular gene expression. We developed a novel CRISPR/Cas9-induced restoration of bioluminescence reporter system, CiRBS, to monitor cellular bioluminescence from a reporter gene in the genome of transgenic *Arabidopsis*. In this method, the enzymatic activity of an inactive *luciferase* mutant*, LUC40Ins26bp*, which has a 26-bp insertion at the 40th codon, was restored by introducing an indel at the insertion site using CRISPR/Cas9. We succeeded in long-term monitoring of the cellular bioluminescence of *Arabidopsis* plants expressing *LUC40Ins26bp*, which was restored by transient transfection with CRISPR/Cas9-inducible constructs using particle bombardment. Recombination events via indels were mostly complete within 24 h of CRISPR/Cas9 induction, and 7.2% of CRISPR/Cas9-transfected cells restored bioluminescence. It was estimated that 94% of the bioluminescence-restored cells carried only one chromosome having the optimal recombination construction. Thus, CiRBS allows for reliable single-cell gene expression analysis of cell-to-cell heterogeneity and temporal fluctuations from a single locus.

## Introduction

Cellular gene expression is the basis of biological systems in multicellular organisms. Gene expression is coordinately regulated but fluctuates stochastically in each cell, and it has been suggested that cellular functions are subject to such noise [1, 2]. However, few studies have been performed on the effects of the stochasticity of individual cells on function at the tissue or organism level. The spatial and temporal analysis of gene expression at the single-cell level within the same organ contributes to the understanding of the role of stochastic fluctuation and cell-to-cell heterogeneity in multicellular organisms.

The circadian clock system of plants is a cell-autonomous system [3–5] composed of cellular oscillators that function via transcription-translation feedback loops of various clock genes [6, 7]. Single-cell analyses have revealed cell type-specific circadian behaviors, tissue-specific synchronization characteristics, and coordination among cellular circadian rhythms in an organ [8–10]. In these studies, the accumulation of the product of a clock gene fused with a fluorescent protein gene was monitored in transgenic *Arabidopsis* plants using confocal microscopy. Given that this reporter system requires an excitation light for fluorescence observation and measurement, it cannot be used to monitor circadian rhythms in darkness. Furthermore, this system is unsuitable for long-term observations with a high temporal resolution of a living plant because of photodamage or unexpected cellular light responses to excitation light [11, 12].

In plants, firefly luciferase (LUC) has been used as a bioluminescent reporter to examine circadian rhythms [13, 14]. In contrast to fluorescent reporters, LUC is an enzyme that emits bioluminescence in cells when a luciferin substrate is added [15], and it is a noninvasive and suitable reporter for the long-term monitoring of gene expression. The bioluminescence rhythms of individual plants, organs, and tissues have been discussed based on the results of transgenic plants expressing *LUC* under the control of circadian promoters, using either a photomultiplier tube or a high-sensitivity camera [16–20]. Monitoring bioluminescent circadian rhythms at the single-cell level in plants has also been reported. Protoplasts of transgenic *Arabidopsis* carrying a *CIRCADIAN CLOCK ASSOCIATED 1 promoter:: luciferase* (*CCA1::LUC*) reporter were observed for their bioluminescence rhythms in culture medium [21]. Furthermore, the cellular circadian behavior in intact plants has been studied using bioluminescence imaging at the single-cell level [22–25].

Using the particle bombardment method, a bioluminescent reporter can be delivered into plant cells with DNA-coated gold particles [26, 27]. Notably, duckweed plants are suitable for long-term bioluminescence monitoring of cells transfected with a reporter using this method. Quantitative single-cell bioluminescence imaging analyses of duckweed plants have demonstrated heterogeneity in clock gene expression among cells within the same organ [22–24, 26]. The particle bombardment method also allows co-transfection of a bioluminescent reporter and effector constructs for overexpression, RNAi, or CRISPR/Cas9-induced genome editing [23, 28–30]. Although this method is suitable for observing circadian behavior in individual cells, comparative analyses of gene expression between cells are limited. The copy number of the reporter gene introduced into the nucleus is heterogeneous among cells due to heterogeneity in the distance between the nuclei and the introduced gold particles [31] and possibly due to heterogeneity in the number of reporter plasmids coating the gold particles. Such heterogeneity results in differences in bioluminescence intensities among transfected cells within the 1000-fold range [26]. Moreover, the bioluminescence intensities of cells transiently transfected with a bioluminescence reporter tend to decrease gradually [22]. These problems can be solved by single-cell analysis of the clock gene expression behavior of a *LUC* reporter gene uniformly introduced into the genome. However, the spatial resolution of the bioluminescence of a stable transgenic plant carrying a *LUC* reporter under the control of a clock gene promoter is insufficient for the single-cell imaging [32, 33]. Therefore, we used a strategy in which cells in a transgenic plant carrying an inactive *LUC* reporter were subjected to genome editing by introducing a *LUC*-reactivating effector via particle bombardment to restore bioluminescence at the single-cell level.

LUC proteins, from various luminous insects, consist of N- and C-terminal domains with a flexible linker sequence between the two [15]. The amino acid residues of the active center to which the luciferin substrate and Mg-ATP bind are highly conserved, and mutated LUCs with high enzymatic activity or different colors have been reported [34–37]. These mutated *LUC*s have been widely used in various bioluminescence imaging experiments [25, 28, 38]. Hence, we hypothesized that we could either mutate *LUC* sequences or insert a few amino acid residues in the LUC protein while keeping the luciferase activity, and use it as the bioluminescence reporter in our reactivating strategy.

We chose the CRISPR/Cas9 system, which consists of clustered regularly interspaced short palindromic repeats (CRISPR) and CRISPR-associated protein 9 (Cas9), as our editing tool [39, 40]. A single guide RNA (*sgRNA*) recognizes and binds to 20-base target sequences, and attracts Cas9, an endonuclease, resulting in double-strand break (DSB) [41]. DSB induces error-prone DNA repair via nonhomologous end-joining (NHEJ) or DNA repair via homology-directed repair (HDR). When repeated sequences exist around the target sequences, HDR occurs immediately after DNA replication and before cell division, and uses homologous double-stranded DNA as template [42]. NHEJ induces point mutations or insertions/deletions (indels) around the DSB site, which alter the functions of the target gene by changing the amino acid sequence, including the introduction of a premature termination codon [43]. A quantitative study on *Arabidopsis* showed that NHEJ-induced mutations were predominantly 1-bp insertions and short deletions [44]. Gene editing by NHEJ is available for various plant species including *Arabidopsis* as a high-efficiency mutagenesis technology [45].

In this study, we developed a <u>C</u>RISPR/Cas9-induced restoration of <u>b</u>ioluminescence reporter <u>s</u>ystem (CiRBS) for single-cell expression analysis that enables cell-to-cell comparison of the gene expression behavior of a luciferase reporter in the genome. Transgenic *Arabidopsis* plants carrying an inactive *LUC* bioluminescence reporter gene were produced, and cellular bioluminescence was restored by particle bombardment transfection with *CRISPR/Cas9* effector constructs to reactivate the *LUC* gene.

A previous study showed that a modified *β-glucuronidase* (*GUS*) reporter (*GU-US*; carrying a repeated sequence around the target sequence of the *sgRNA*) expressed in a transgenic potato plant was reactivated through HDR by *Agrobacterium*-mediated transfection with *CRISPR/Cas9* constructs [46]. This reactivation treatment resulted in 2-fold increases in reporter activity compared to that in the negative controls. In our study, we investigated the conditions under which the modified *LUC* exhibited null activity, and such activity was restored by indels through NHEJ. We first identified a site in the *LUC* gene at which a 24-bp sequence insertion, including the *sgRNA* target, retained the luciferase activity of the gene product. We then constructed a modified *LUC* with a 26-bp insertion at the same site to inactivate luciferase activity by frameshift and stop codon. This 26-bp insertion also included the *sgRNA* target sequence. Finally, transgenic *Arabidopsis* plants carrying this construct were created. The modified *LUC* was reactivated by transfection with *CRISPR/Cas9* constructs using particle bombardment. Hence, we successfully monitored the bioluminescence of the reporter gene in the genome at the single-cell level.

## Results

### Design of six insertion sites in the *LUC* gene for CRISPR/Cas9 target sequences

We generated mutant *LUC* constructs with specific 24-bp or 26-bp insertions for the CiRBS (Fig. 1a). *LUCX_AA_Ins24bp* and *LUCX_AA_Ins26bp* included 24-bp and 26-bp insertions after the X_AA_th codon from the start codon (ATG) in the *LUC* gene, respectively (Fig. 1b). The first 20-bp were common to every construct, and included the *sgRNA* target sequence for the CRISPR/Cas9 system, as previously reported [46]; no sequence matching this 20-bp sequence was found in the *Arabidopsis* genome. The same PAM sequence (CGG) was used for all target sequences. *LUCX_AA_Ins26bp* insertions resulted in *LUC* deletion mutants due to the formation of a stop codon and a frameshift at the end of each insertion (Fig. 1c). The third nucleotide (A) of the stop codon (TAA) belonged to the original *LUC* gene. *LUCX_AA_Ins26bp* insertions were expected to generate null mutants, whose bioluminescence activity could be restored by CRISPR/Cas9-induced indels in the insertion sequences themselves. *LUCX_AA_Ins24bp* represented revertant *LUC* mutants. LUCX_AA_Ins24bp proteins included an 8-AA sequence (V-L-P-M-I-P-S-G) at the insertion site. We selected six insertion sites after codons 2nd, 40th, 106th, 239th, 378th, and 491st from the *LUC* start codon.

**Fig. 1.**
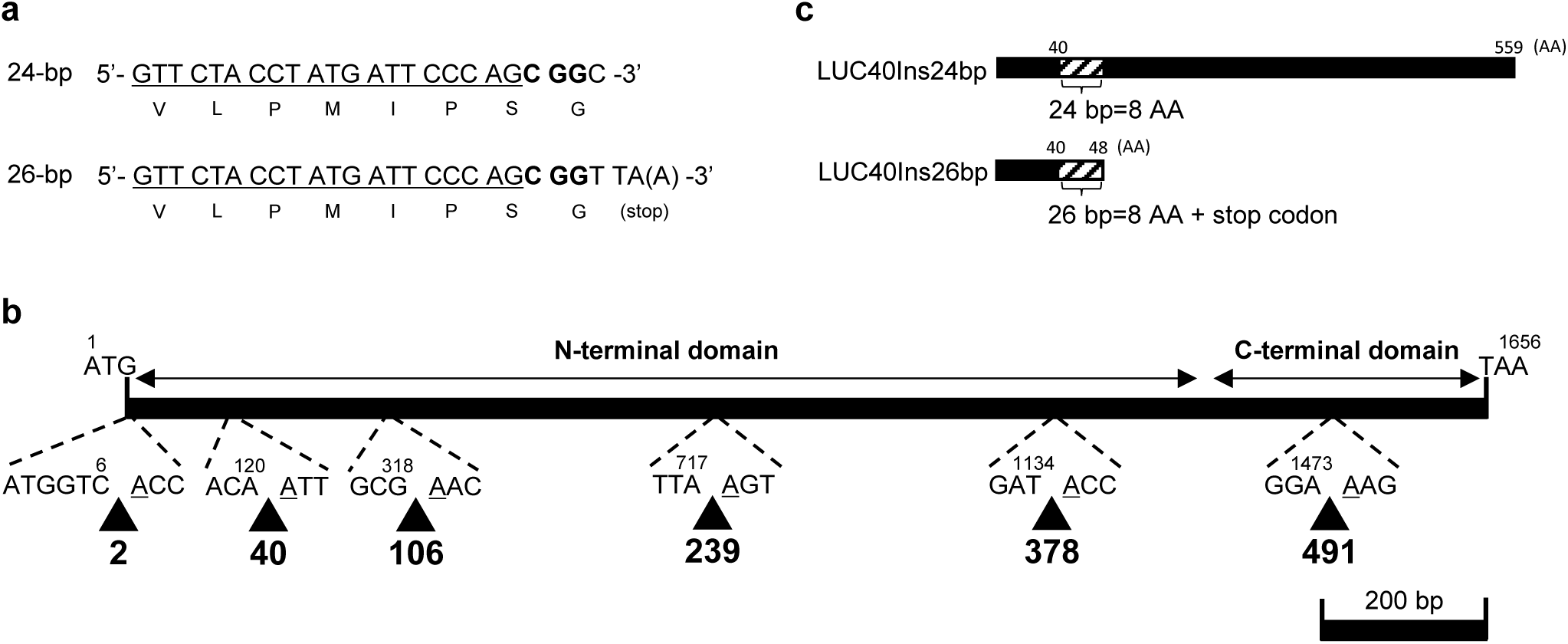
Schematic representation of *LUCX_AA_Ins24bp* and *LUCX_AA_Ins26bp* inserts. (**a**) Nucleotide and amino acid (translated below each codon) sequences of the insertion in *LUCX_AA_Ins24bp* (top) and *LUCX_AA_Ins26bp* (bottom). The target sequences of the *sgRNA* of the CRISPR/Cas9 system are underlined. Characters in bold indicate the PAM sequence CGG. The adenine in parentheses in the *LUCX_AA_Ins26bp* sequence represents the first nucleotide of the *LUC* gene, which is involved in the generation of a stop codon at the insertion site. (**b**) Map of insertion sites. Arrowheads indicate the insertion sites for *LUCX_AA_Ins24bp* and *LUCX_AA_Ins26bp*. The number below each arrow represents the codon (X_AA_) next to which either *LUCX_AA_Ins24bp* or *LUCX_AA_Ins26bp* were inserted. The N- and C-terminal domain coding regions are represented by horizontal lines [15]. (**c**) Schematic representation of the proteins encoded by *LUCX_AA_Ins24bp* (top) and *LUCX_AA_Ins26bp* (bottom) inserted at 40th codon (LUC40Ins24bp and LUC40Ins26bp, respectively). Shaded boxes represent the 8-AA inserted regions.

### Selection of the *LUC40Ins26bp* for CiRBS

To determine which LUCX_AA_Ins24bp proteins retained the highest bioluminescence activity, we quantified the bioluminescence intensity of each *LUCX_AA_Ins24bp* variant. We transiently transfected duckweed (*Lemna japonica*) plant cells with a *CaMV35S::LUCX_AA_Ins24bp* construct using particle bombardment. *CaMV35S::LUCX_AA_Ins24bp* contains a *LUCX_AA_Ins24bp* coding sequence under the control of the *Cauliflower mosaic virus 35S* (*CaMV35S*) promoter. Duckweed plants are suitable materials for bioluminescence imaging at the single-cell level because of their tiny flat bodies [22, 26, 33]. Bioluminescence imaging was performed one day after gene transfection (Fig. 2a). Bioluminescent spots were observed in all *CaMV35S::LUCX_AA_Ins24bp* transfected cells except for those with *CaMV35S::LUC239Ins24bp*. We then checked for the loss of bioluminescence activity in each *CaMV35S::LUCX_AA_Ins26bp* construct (Fig. 2b). Almost no bioluminescent spots were observed for any of the *CaMV35S::LUCX_AA_Ins26bp*-expressing cells except for those with *CaMV35S::LUC2Ins26bp*.

**Fig. 2.**
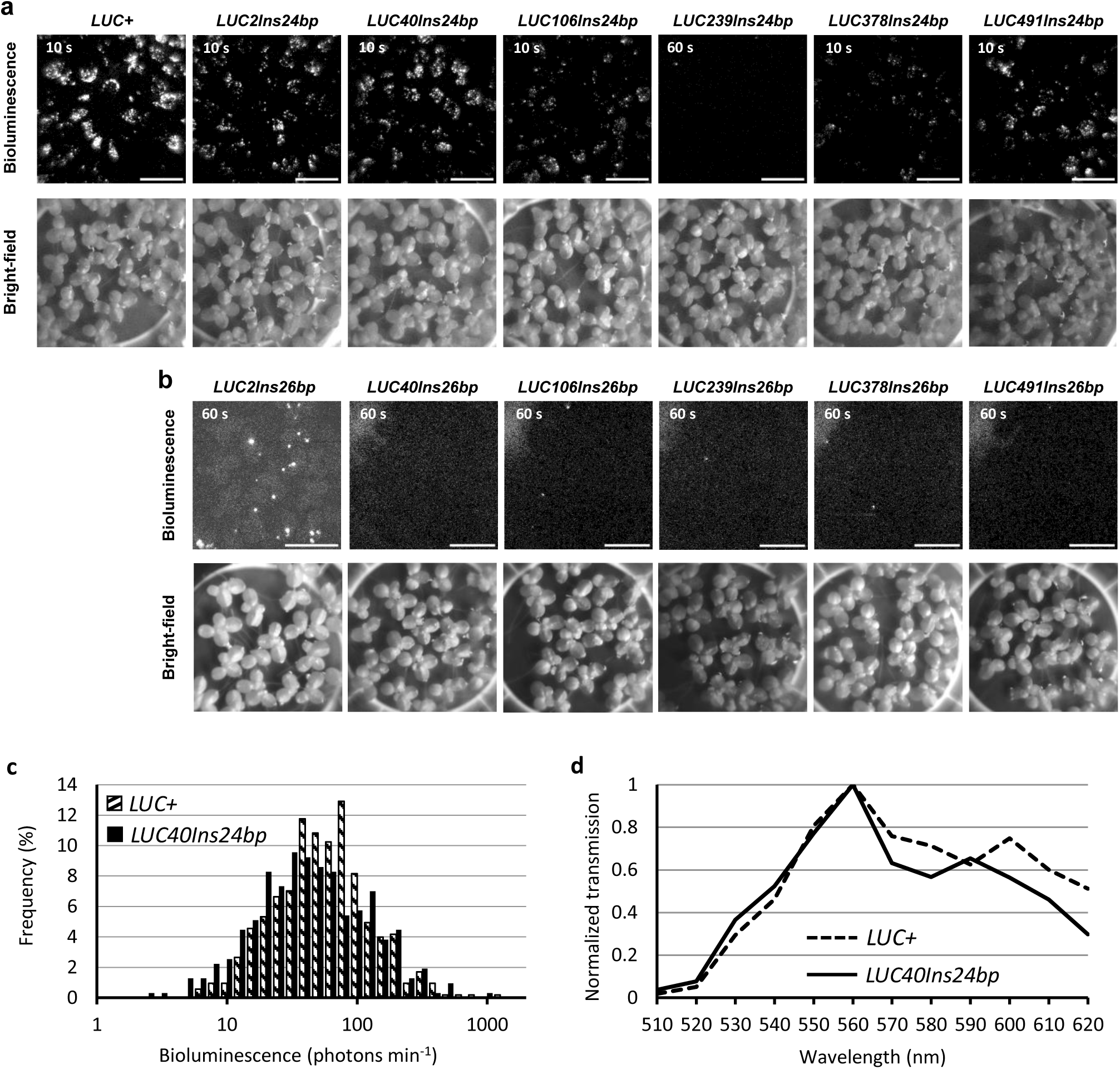
Bioluminescence of *LUCX_AA_Ins24bp* and *LUCX_AA_Ins26bp*. (**a**, **b**) Bioluminescence (top) and bright-field (bottom) images of duckweed plant cells transfected with (**a**) *CaMV35S::LUCX_AA_Ins24bp* or (**b**) *CaMV35S::LUCX_AA_Ins26bp*. The exposure time for capturing bioluminescence is shown in each image. The signal range for the bioluminescence images was fixed to (**a**) 1920–5000 or (**b**) 1920–2500. The signal range of raw 16-bit images was 0–65535, whereas that of the background was 1920. Bars: 10 mm. (**c**) Frequency distribution (%) of cellular bioluminescence intensities for *CaMV35S::LUC+* (n = 526) and *CaMV35S::LUC40Ins24bp* (n = 314). (**d**) Bioluminescence spectra of LUC+ and LUC40Ins24bp in duckweed plants. The transmitted bioluminescence intensities were normalized to that at 560 nm and plotted. Bioluminescence was filtered with a series of band-pass filters and quantified.

We quantified the bioluminescence intensities of individual spots for each *CaMV35S::LUCX_AA_Ins24bp*- and *CaMV35S::LUC+*-expressing cells (Fig. 2c; Supplementary Fig. S1). At this time, the bioluminescence of *CaMV35S::LUC40Ins24bp* was higher than that of any other *CaMV35S::LUCX_AA_Ins24bp* constructs, and the bioluminescence intensity distribution in *CaMV35S::LUC40Ins24bp* and *CaMV35S::LUC+* cells was similar (Fig. 2c). We also examined the emission spectra of *CaMV35S::LUCX_AA_Ins24bp* and *CaMV35S::LUC+* and confirmed that those of *LUC40Ins24bp* and *LUC+* were similar (Fig. 2d; Supplementary Fig. S2). We concluded that the insertion at the 40th codon was the best among the six candidate sites, and used *LUC40Ins26bp* for further CiRBS experiments.

We then examined NHEJ-induced restoration of bioluminescence of *LUC40Ins26bp*. In order to test the effectiveness of our *sgRNA*, duckweed cells were transiently co-transfected with both *CaMV35S::LUC40Ins24bp* and *CRISPR/Cas9* constructs by particle bombardment. *CRISPR/Cas9* constructs consisted of a mixture of *pENTR_AtU6-26::sgRNA-LUCX_AA_* and *pUC18_PcUBQ4-2::Cas9* [47]. The cellular bioluminescence intensities of *CaMV35S::LUC40Ins24bp* in the co-transfected samples were considerably lower than in those carrying *CaMV35S::LUC40Ins24bp* only (Fig. 3a-3c). Furthermore, co-transfection with a nontargeting mixture containing *pENTR_AtU6-26::sgRNA* and *pUC18_PcUBQ4-2::Cas9* (nonsense-*CRISPR/Cas9* constructs) did not lead to decreased bioluminescence intensities (Supplementary Fig. S3). We then transiently co-transfected duckweed cells with *CaMV35S::LUC40Ins26bp* and the *CRISPR/Cas9* constructs. The number of bioluminescent spots dramatically increased (Fig. 3a), whereas few bioluminescent spots were observed for cells transfected with *CaMV35S::LUC40Ins26bp* without the *CRISPR/Cas9* constructs (Fig. 2b; Fig. 3a). Given that NHEJ-induced mutations are predominantly 1-bp insertions and short deletions [44], luciferase activity restoration was likely triggered by these indels at the insertion site of *LUC40Ins26bp*. The geometric mean of bioluminescence intensities of restored-*LUC40Ins26bp* was 3% of that of *LUC40Ins24bp* (Fig. 3c). This suggests that a small portion of the introduced *LUC40Ins26bp* copies by particle bombardment was restored through NHEJ. We also transiently co-transfected duckweed cells with both *CaMV35S::LUC40Ins26bp* and the nonsense-*CRISPR/Cas9* constructs, but no bioluminescent spots were observed (Supplementary Fig. S3).

**Fig. 3.**
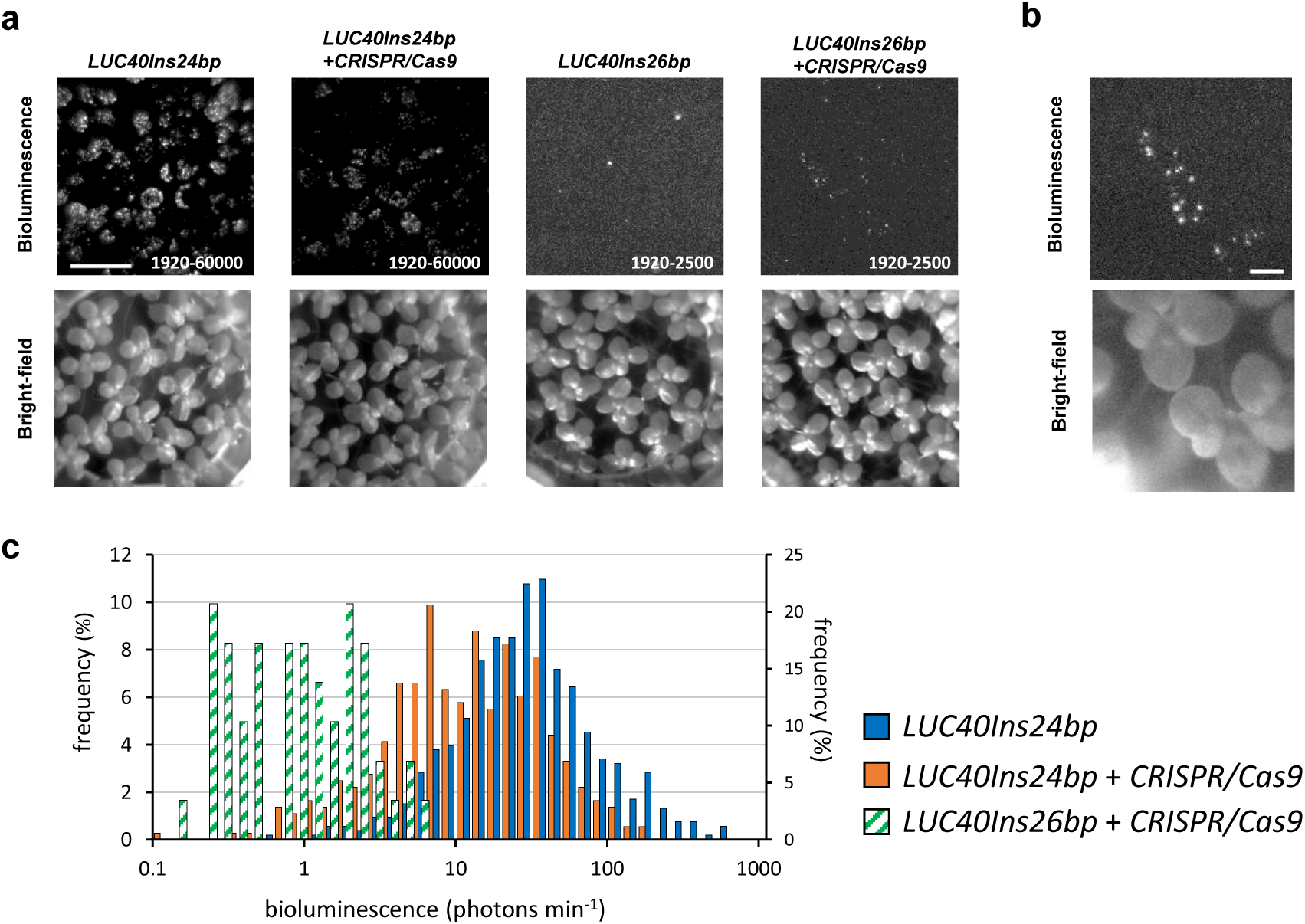
Restoration of *LUC40Ins26bp* bioluminescence by co-transfection with *CRISPR/Cas9* constructs. (**a**) Bioluminescence (top) and bright-field (bottom) images of duckweed plants transfected with reporter and *CRISPR/Cas9* constructs (indicated above each set of images). Signal ranges are shown in each bioluminescence image. Exposure time: 60 s; Bar: 10 mm. (**b**) Close-up images of duckweed plants co-transfected with *CaMV35S::LUC40Ins26bp* and *CRISPR/Cas9* constructs. Exposure time: 180 s. The signal range: 1920–2500. Bar: 2 mm. (**c**) Frequency distribution (%) of cellular bioluminescence intensities for *CaMV35S::LUC40Ins24bp* (blue bars; n = 529; left side *y*-axis), *CaMV35S::LUC40Ins24bp* and *CRISPR/Cas9* constructs (orange bars; n = 364; left side *y*-axis), and *CaMV35S::LUC40Ins26bp* and *CRISPR/Cas9* constructs (green shaded bars; n = 29; right side *y*-axis).

We further tested the bioluminescence restoration of *LUC40Ins26bp* using a plant-derived promoter commonly employed in physiological experiments. We chose a circadian promoter of the *Arabidopsis* clock gene, *CIRCADIAN CLOCK ASSOCIATED 1* (*AtCCA1*) [4]. Duckweed plants transfected with *AtCCA1::LUC+* or *AtCCA1::LUC40Ins24bp* via particle bombardment showed a bioluminescence circadian rhythm [48] (Supplementary Fig. S4). In contrast, the bioluminescence of plants transfected with *AtCCA1::LUC40Ins26bp* did not show a circadian rhythm, and its intensities were close to the background level. Notably, co-transfection with *AtCCA1::LUC40Ins26bp* and *CRISPR/Cas9* constructs restored bioluminescence and exhibited circadian rhythms similar to those of *AtCCA1::LUC+* and *AtCCA1::LUC40Ins24bp*, although with lower intensities. Therefore, the *LUC40Ins26bp* reporter system is likely to function without altering the physiological relevance of the promoter.

### Cellular bioluminescence restoration in transgenic *Arabidopsis* carrying *CaMV35S::LUC40Ins26bp*

We tested CiRBS in transgenic *Arabidopsis* plants carrying *CaMV35S::LUC40Ins26bp* integrated in the genome. We selected a T3 homozygous line, *LUC40Ins26bp #28-3*, which strongly expressed *LUC40Ins26bp* from a single locus (Supplementary Fig. S5). We transiently transfected detached leaves of the transgenic line with *CRISPR/Cas9* constructs using particle bombardment. Immediately after transfection, the leaves were floated on a luciferin-containing medium and monitored for cellular bioluminescence under constant dark conditions for one week. We successfully monitored restored bioluminescence at the single-cell level (Fig. 4a). This restored bioluminescence appeared to be stably maintained when compared to that in *Col-0* detached leaves with conventional transient transfection with *CaMV35S::LUC+* (Fig. 4b). Few bioluminescent spots were detected when nonsense-*CRISPR/Cas9* constructs were used for transfection, and no bioluminescent spots when only *pENTR_AtU6-26::sgRNA-LUCX_AA_* or only *pUC18_PcUBQ4-2::Cas9* were used.

**Fig. 4.**
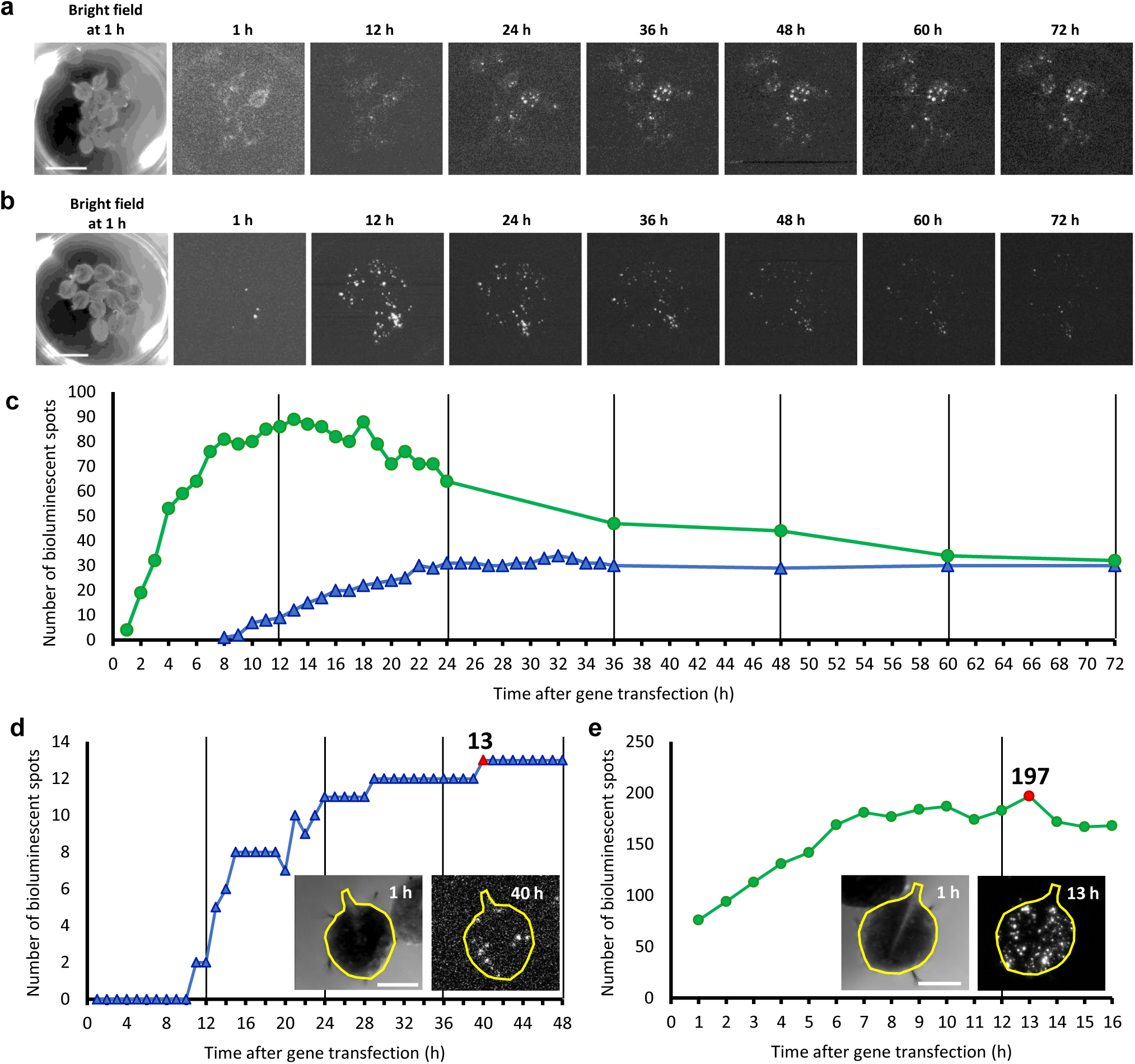
Time-course of the bioluminescence restoration events in transgenic *Arabidopsis* carrying *CaMV35S::LUC40Ins26bp* (line *#28-3*). (**a, b**) Low-magnification bioluminescence images of the (**a**) bioluminescence-restored *CaMV35S::LUC40Ins26bp* and (**b**) transiently-transfected *CaMV35S::LUC+ Arabidopsis* leaves. Time 0 represents the image capturing starting time, immediately after gene transfection. *Arabidopsis* leaves were recorded under constant dark conditions. Exposure time: (**a**) 150 and (**b**) 60 s. The signal range was fixed to 1920–2200. Bar: 10 mm. (**c**) Number of bioluminescent spots with over 2000 signals in (**a)** (blue △) and (**b)** (green 〇). (**d, e**) Number of bioluminescent spots in (**d**) restored-bioluminescence *CaMV35S::LUC40Ins26bp* and (**e**) transiently-transfected *CaMV35S::LUC+ Arabidopsis* leaves. The bioluminescent spots in the leaf with the highest gene transfection efficiency for each treatment were counted. Red symbol: maximum number of bioluminescent spots. The bright-field (left) and bioluminescence (right) images of the corresponding leaves (together with a representation of the shape of each leaf) are shown. Bioluminescence monitoring conditions were the same as in (**a**, **b**) except for the magnification used for a better visualization of the spots. The areas of the leaves are (**d**) 27,045 and (**e**) 29,568 pixel^2^. Bars: 2.5 mm.

To quantitatively analyze the timing of bioluminescence emergence, we counted the bioluminescent spots in a series of time-lapse images for Fig. 4a and 4b. These bioluminescent spots were detected immediately after *CaMV35S::LUC+* transient transfection of *Col-0* detached leaves (Fig. 4b and 4c). The number of spots increased after 12 h, and then gradually decreased. The transiently transfected reporter appeared to be unstable in cells. In contrast, when detached leaves of the *LUC40Ins26bp #28-3* line were transfected with the *CRISPR/Cas9* constructs, we first detected a restored bioluminescent spot at 8 h, and the number of spots gradually increased for up to 24 h (Fig. 4a and 4c). Thereafter, the number was maintained, and bioluminescence appeared to persist. This indicated that most bioluminescence restoration events were completed within 24 h of gene transfection with *CRISPR/Cas9*, and cellular bioluminescence was stably maintained. In these leaves, it took a longer time for bioluminescent spots to emerge compared to those observed after transient transfection with *CaMV35S::LUC+*, probably because of the multiple molecular events required for such restoration, including the expression of *CRISPR/Cas9* constructs, DSB, NHEJ, and recombinant *LUC40Ins26bp* expression. Although some bioluminescent spots emerged several days after *CRISPR/Cas9* construct transfection, bioluminescence was basically restored the day after transfection.

We tried to estimate the efficiency of bioluminescence restoration by NHEJ in the transfected cells, and the apparent efficiency was defined as the ratio of the number of restored bioluminescent spots to the number of transiently transfected cells in *LUC40Ins26bp #28-3* leaves. However, we were unable to simultaneously detect these two cell types in the same leaf. To estimate the ratio, we counted the maximum number of bioluminescent spots in a leaf of wild-type transfected with *CaMV35S::LUC+*, and also counted the maximum number of restored bioluminescent spots in a leaf of the *LUC40Ins26bp #28-3* line transfected with the *CRISPR/Cas9* constructs (Fig. 4d and 4e). The number of bioluminescent spots was 197 at 13 h for wild-type leaves (Fig. 4d), and 13 at 40 h for *LUC40Ins26bp #28-3* leaves (Fig. 4e). Based on these numbers, we estimated the apparent efficiency as follows:

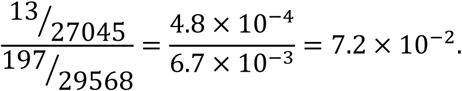

The number of bioluminescent spots was corrected for leaf size. Thus, it was presumed that 7.2% of total transfected cells restored the luciferase activity in our experimental conditions.

We next estimated the probability of a recombination event that restored luciferase activity at the *LUC40Ins26bp* locus in each chromosome. In *Arabidopsis* leaves, the ploidy levels of cells range from 2C to 32C owing to endoreduplication [49, 50]. The reported ratio in epidermal cells is 2n:4n:8n:16n:32n = 0.32:0.38:0.19:0.10:0.01 [49]. In this study, we referred to this ratio to calculate the probability of recombination events that restore bioluminescence. Given our estimation that bioluminescence-restored cells accounted for 7.2% of the cells transfected with *CRISPR/Cas9* constructs, the optimal recombination rate (α) at the *LUC40Ins26bp* locus was 1.37% (Supplementary Fig. S6). It was also calculated that 94% of the bioluminescence-restored cells carried only one chromosome with the optimal recombination (Fig. 5). Therefore, the bioluminescence intensities of most cells were presumed to reflect the expression level of the *LUC40Ins26bp* locus on a single chromosome in the single-cell CiRBS.

**Fig. 5.**
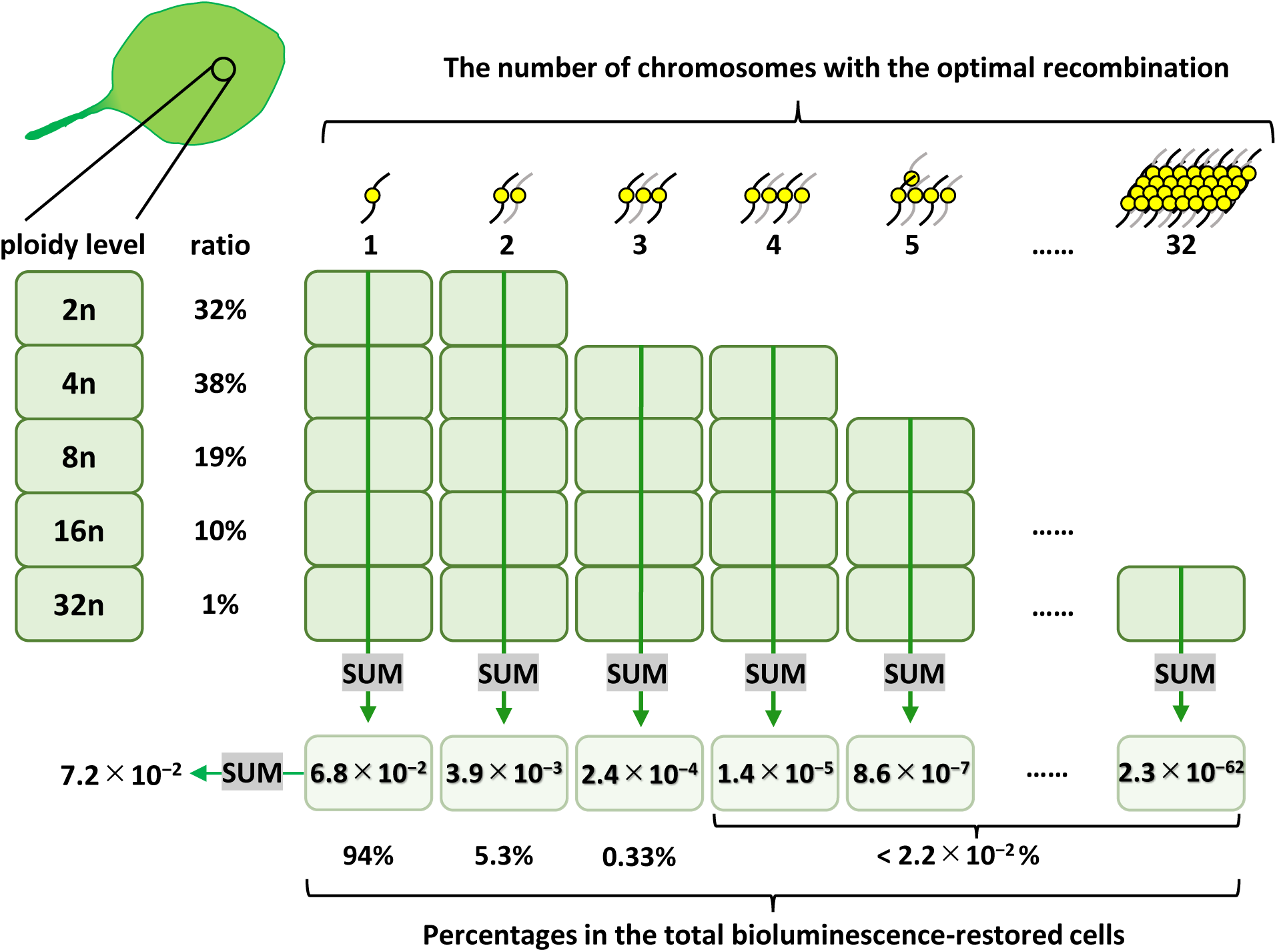
Estimation of the ratios of transfected cells carrying various number of chromosomes with the optimal recombination for bioluminescence restoration. We referred to the ploidy profile of *Arabidopsis* leaf epidermal cells (2n:4n:8n:16n:32n = 0.32:0.38:0.19:0.10:0.01) reported by Kawade and Tsukaya, 2017 [49]. Based on a Poisson distribution with the probability of the optimal recombination (α=0.0137; see Supplementary Fig. S6), we calculated the ratios of bioluminescence-restored cells to transfected cells in every combination of chromosome number and ploidy level. The ratios of bioluminescence-restored cells carrying 1n–32n chromosomes with the optimal recombination are represented. The percentages of cells carrying 1, 2, 3, and > 4 chromosomes with the optimal recombination among all bioluminescence-restored cells are also represented.

### Observation of cellular bioluminescence traces by CiRBS

To demonstrate the single-cell analysis using CiRBS, we transiently transfected cells into the detached leaves of *LUC40Ins26bp #28-3* with *CRISPR/Cas9* constructs using particle bombardment, and monitored the restored bioluminescence of *LUC40Ins26bp* for a week. Leaves were collected from plants grown under 12-h light/12-h dark conditions before gene transfection (Fig. 6a). We quantified the bioluminescence intensities of the seven spots whose bioluminescence continued during the measurements (Fig. 6b and 6c). The bioluminescence of each spot appeared to fluctuate asynchronously in the same detached leaf, suggesting that fluctuations in gene expression in the genome of individual cells can be analyzed using CiRBS. The mean bioluminescence intensities of the seven spots remained nearly constant under constant light. In another strain (*LUC40Ins26bp #38-2*), we observed cellular bioluminescent traces with a seemingly circadian fluctuation as well as those with asynchronous fluctuations (Supplementary Fig. S7). Intensities were comparable within a 10-fold range at each time point. Interestingly, it has been reported that transient transfection with *CaMV35S::LUC+* results in a large variation in bioluminescence intensity, over a 1000-fold range, among cells [26]. These results suggest that the observation of cellular bioluminescence by CiRBS allows us to compare cellular gene expression behavior, including expression levels and stochastic fluctuations between cells.

**Fig. 6.**
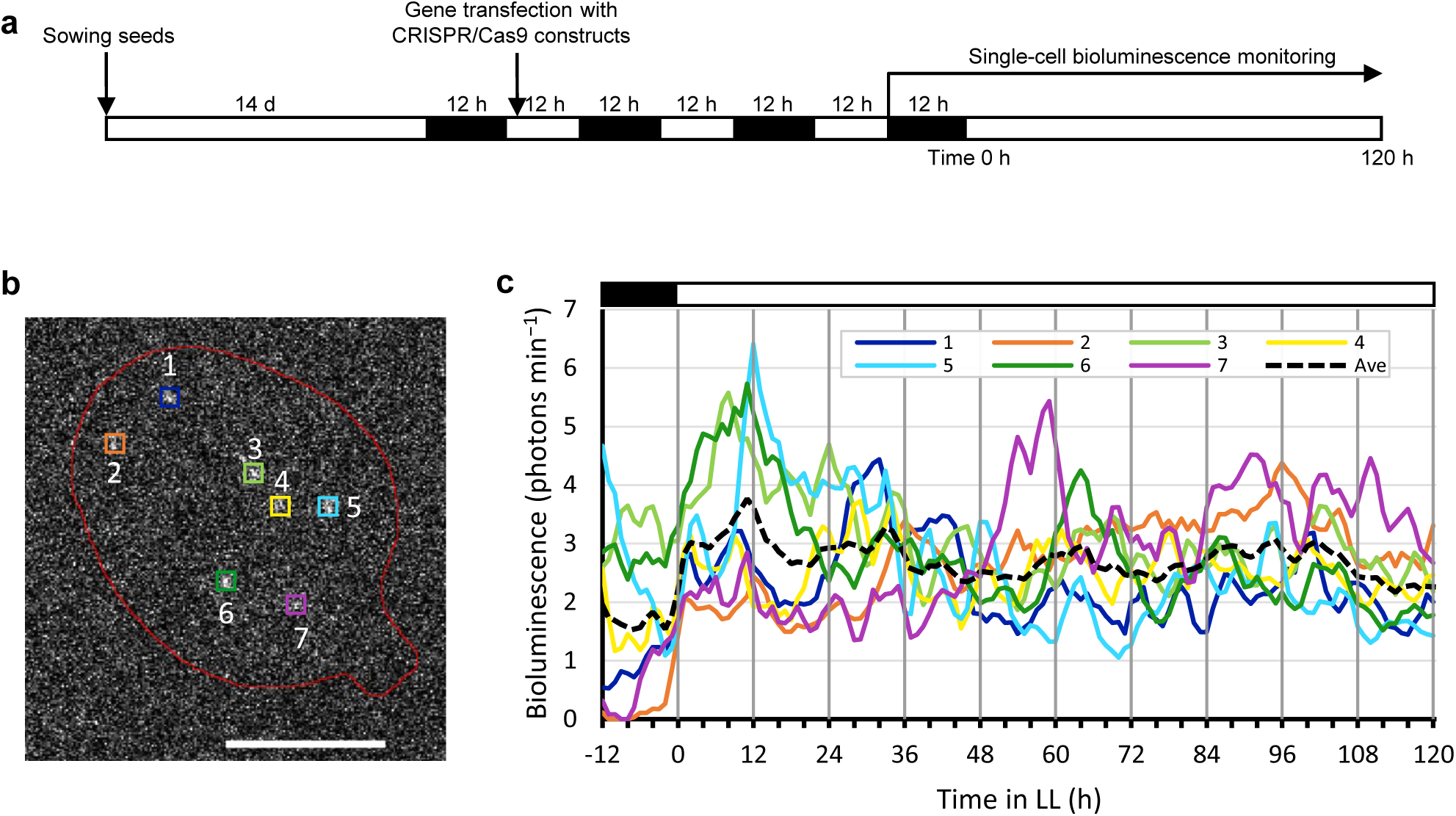
Bioluminescence traces of restored-*LUC40Ins26bp* at the single-cell level. (**a**) Time schedule of the monitoring experiments. Gene transfection of cells in detached leaves with *CRISPR/Cas9* constructs was conducted by particle bombardment. Two days after gene transfection, the leaf showing the largest number of bioluminescent spots was selected for bioluminescence monitoring. Time 0 indicates the start of the constant light. Hourly bioluminescence imaging was performed during the monitoring. White and black boxes represent light and dark conditions, respectively. (**b**) Snapshot of bioluminescent spots (colored □) at 3 h. The seven bioluminescent spots of interest were numbered from the distal to the proximal part of the leaf. The shape of the leaf is roughly represented. Bar: 2 mm. (**c**) Bioluminescence traces of the seven spots. Quantified intensities (3-h moving average) are plotted for each bioluminescent spot shown in (**b**). Mean intensities for the seven spots are represented by a black dotted line.

## Discussion

We successfully developed a CiRBS, which enabled gene expression analysis at the single-cell level, and demonstrated fluctuations in *luciferase* gene expression driven by the *CaMV35S* promoter on the genome of individual cells. This CiRBS is based on the NHEJ of the CRISPR/Cas9 system to restore the luciferase activity of a loss-of-function *LUC* mutant. Previous reports have shown gene restoration through HDR triggered by CRISPR/Cas9 between repeated sequences [46, 51]. Therefore, we first attempted this strategy and produced a modified *LUC* gene with 702-bp repeated sequences before and after the *sgRNA* target sequence in the coding region (named *LU_UC* gene; Supplementary Fig. S8). Although *LU_UC* was expected to be a complete loss-of-function gene, transient transfection of leaves with *CaMV35S::LU_UC* resulted in a much stronger bioluminescence emission than the background signal of the sample. The retained luciferase activity of *LU_UC* restricted its usage in the CiRBS. In addition, transient co-transfection with *LU_UC* and *CRISPR/Cas9* constructs did not considerably increase the bioluminescence intensity (Supplementary Fig. 9). Considering that HDR is more frequently induced during cell division [42], NHEJ is likely to be more dominant than HDR in mature cells, which are the targets of transfection by particle bombardment [52].

We then aimed to develop a simple reporter gene with little or no bioluminescence background. When comparing the LUC sequences of various luminous insects, the active sites were found to be highly conserved [15]. To construct CiRBS using NHEJ, we investigated the sites at which a short amino acid insertion would not affect luciferase activity. *LUCX_AA_Ins24bp* construction, which included a 24-bp insertion sequence (itself including the *sgRNA* target sequence) inserted at certain X_AA_ sites of the *LUC+* gene, showed various enzymatic activities depending on the X_AA_ insertion site (Fig. 2a and 2c; Supplementary Fig. S1). LUC2Ins24bp and LUC40Ins24bp showed bioluminescence intensities comparable to those of wild-type luciferase LUC*+*. This suggests that luciferase is robust against mutations at these sites. In contrast, LUC239Ins24bp was a mutant LUC protein in which the 24-bp sequence was inserted near the active site, resulting in activity loss. Additionally, it has been reported that the structure around the active site is related to the color of LUC bioluminescence [53]. Therefore, we examined the bioluminescence spectra of LUCX_AA_Ins24bp proteins. While LUC106Ins24bp exhibited a strong red emission (>590 nm), all other LUCX_AA_Ins24bp did not considerably change their spectra (Supplementary Fig. S2; Fig. 2d).

By introducing both a frameshift and a stop codon (TAA) at the end of the 26-bp insertion, we succeeded in losing the luciferase activity of LUCX_AA_Ins26bp proteins, except for that of LUC2Ins26bp (Fig. 2b). We expected that LUC2Ins26bp would lose its activity because it only encodes a 10-AA peptide sequence from the start codon of the native protein. Its luciferase activity might be caused by an N-terminal 30-AA deleted mutant LUC, starting from an internal in-frame Met codon. It should be noted that LUC491Ins26bp lost its enzymatic activity, although most of the LUC sequence remained intact (Fig. 1; Fig. 2). This finding supports a previous report indicating that the C-terminal domain of luciferase plays a crucial role in the stable binding of luciferin and the subsequent luminescence reaction [53].

We selected *LUC40Ins26bp* as the reporter gene for CiRBS because the 24-bp insertion maintained luciferase activity, whereas the 26-bp insertion lost it (Fig. 2a and 2b). The amplitudes of the bioluminescence rhythms of *AtCCA1::LUC+* and *AtCCA1::LUC40Ins24bp* were similar (Supplementary Fig. S4), suggesting that the stability of LUC40Ins24bp was similar to that of LUC+ [54]. However, we detected a few bioluminescent spots on a small number of fronds (leaf-like structures) when duckweed plants were transfected with *CaMV35S::LUC40Ins26bp* (Fig. 3a). We tested the possibility that an alternative translation start site located after the insertion (cLUC40) could be utilized to produce a partially deleted luciferase protein that retained its enzymatic activity when in the presence of the N-terminal polypeptide stopped at the insertion site (nLUC40) (Supplementary Fig. S10). However, co-transfection with two genes encoding these polypeptides did not restore bioluminescence activity. Alternatively, cells with a reversion mutation in the insertion region may have partially regained luciferase activity. However, the luminescence of these bioluminescent spots tended to dampen quickly, suggesting that a stable mutation was unlikely. Transcription errors that induce reversion of luciferase activity may occur with a low probability [55]. Therefore, these transcriptional errors may have caused the low background luminescence signals observed after adding luciferin to the medium (Fig. 4a). Notably, the background and stochastic luminescence rarely affected the performance of single-cell CiRBS monitoring and analysis.

Based on our data regarding the emergence of bioluminescence (Fig. 4), we estimated the time required for recombination processes induced by transfection with the *CRISPR/Cas9* constructs using particle bombardment. After the cells in *Col-0* leaves were transiently transfected with *CaMV35S::LUC*+, bioluminescent spots were detected within 1 h, suggesting that the expression of genes transfected via particle bombardment occurred rapidly. In contrast, it was previously reported that indel mutations were first observed 6 hours after introducing CRISPR/Cas9 into tomato protoplasts and their number increase for up to 2–3 d [56]. Consistent with this report, it took 8 h to detect the first restoration of bioluminescence after gene transfection using our CiRBS. Most bioluminescent spots emerged within 24 h, and a small number of bioluminescent spots appeared several days after gene transfection. Various events, including DSB, NHEJ, and transcription-translation of recombinant genes, likely contributed to the delay in bioluminescence emergence.

In this study, we successfully detected cellular bioluminescence from a luciferase driven by the constitutive promoter *CaMV35S* as a model for the single-cell analysis using CiRBS (Fig. 6). Seven spots observed in transgenic *Arabidopsis* (*LUC40Ins26bp #28-3*) showed fluctuations without circadian rhythmicity. This contrasts with our previous observations in duckweed plants transfected with *CaMV35S::LUC+*, which showed circadian rhythms under constant light [25, 28, 48]. Interestingly, bioluminescence rhythms were observed in a portion of the bioluminescent spots in another transgenic *Arabidopsis* line (*LUC40Ins26bp #38-2*) (Supplementary Fig. S7). These results suggested that the differences in circadian rhythmicity were due to positional effects of the transgene, and that the circadian bioluminescence rhythm was stochastically generated in each cell [28, 57]. Cellular bioluminescence intensity was comparable within a 10-fold range of the mean value. In *Arabidopsis* leaves, the ploidy levels of cells range from 2C to 32C owing to endoreduplication [49, 50]. The variation in cellular bioluminescence intensities might be partly due to differences in the number of restored *luciferase* genes among the cells. However, we estimated that 94% of the bioluminescence-restored cells carried only one chromosome with the optimal recombination (Fig. 5); the bioluminescence intensities of most cells reflected the expression level of a single reporter locus in one chromosome. Consequently, single-cell bioluminescence intensity comparison analyses using CiRBS will be highly reliable.

Observation of cellular bioluminescence by CiRBS allows us to compare cellular gene expression behavior, including expression levels and stochastic fluctuations between cells. In other words, the CiRBS at a single-cell level will make it possible to simultaneously capture “fluctuations in expression (temporal changes within the single-cell)” and “heterogeneity (differences between cells),” which are currently difficult to analyze simultaneously.

## Methods

### Plant material and growth conditions

Duckweed plants (*Lemna japonica 5512*) were maintained in Non-flowering (NF) medium with 1% sucrose at 25 ± 1 °C under constant light conditions as previously described [25]. White light (30–35 µmol m^−2^·s^−1^) was supplied by fluorescent lamps (FLR40SEX-W/M/36-HG; NEC, Tokyo, Japan). *L. japonica* plants were grown on 60–80 mL of NF medium in 300-mL Erlenmeyer flasks plugged with cotton to prevent contamination. New stock cultures were prepared once every two weeks, and well-grown plants were used for the experiments.

*Arabidopsis thaliana* plants (*Col-0* and transgenic *CaMV35S::LUC40Ins26bp*/*Col-0*) were grown in plates containing 0.5× Murashige and Skoog (MS) medium with 0.8% agar and 1% sucrose at 22 °C under constant light conditions as previously described [21]. White light (30–35 µmol m^−2^·s^−1^) was supplied by fluorescent lamps (FLR40SEX-W/M/36-HG; NEC). We used 14-day-old *Arabidopsis thaliana* plants and cut the first and second leaves with sterile scissors for particle bombardment. *Arabidopsis* seeds were sterilized using the following procedure: seeds were collected in sterilized 1.5 mL Eppendorf tubes and washed once with 70% ethanol, the supernatant was removed, a seed sterilization solution (5% bleach [Kao, Tokyo, Japan] and 0.05% Triton x-100 [nacalai tesque, Kyoto, Japan]) was added, and the tubes were shaken for 10 min. The seeds were then washed three times with sterile water, and 1 mL of sterile water was added to the tubes, which were shaded at 4 °C for 2 d before culturing on MS medium. Seed sterilization for the selection of transgenic plants was performed using the chlorine gas method (10 mL HCl to 100 mL 5% bleach) [58]. The seeds were sterilized in a sealed desiccator for 6 h.

### Gene transfection constructions

*pUC18_CaMV35S::LUCX_AA_Ins24bp* and *pUC18_CaMV35S::LUCX_AA_Ins26bp* were constructed as follows: We first constructed *pENTR_LUC+* by inserting the luciferase coding sequence (LUC+; Promega, Madison, WI, USA) into the cloning site of the *pENTR/D-TOPO* vector (Thermo Fisher Scientific, Waltham, MA, USA). Then, we constructed *pENTR_LUCX_AA_Ins24bp/26bp*, a derivative of *pENTR_LUC+*.

*pENTR_LUC2Ins24bp/26bp* were generated using a KOD-Plus-Mutagenesis Kit (TOYOBO, Osaka, Japan) with a set of specific primers (Supplementary Table S1). Circularization was performed using the In-fusion method (In-Fusion® Snap Assembly Master Mix; Takara Bio, Shiga, Japan). To construct *pENTR_LUC40Ins24bp/26bp*, *pENTR_LUC106Ins24bp/26bp*, *pENTR_LUC239Ins24bp/26bp*, *pENTR_LUC378Ins24bp/26bp*, and *pENTR_LUC491Ins24bp/26bp*, we prepared three DNA fragments: *5’-part-LUC*, *3’-part-LUC*, and *pENTR/Not*I*_Asc*I. The *5’-part-LUC* fragment for each construct consisted of a PCR amplification product from the start codon to the insertion site of the *LUC* gene in *pENTR_LUC+*. A common forward primer (5’-GCAGGCTCCGCGGCCGCCATG-3’) and a reverse primer specific for each construct were used for amplification (Supplementary Table S2). Each *3’-part-LUC* fragment was a PCR amplification product from the insertion site to the stop codon of the *LUC* gene in *pENTR_LUC+*. A common reverse primer (5’-AAGCTGGGTCGGCGCGCCTTA-3’) and a forward primer specific for each construct were used for the PCR amplification (Supplementary Table S2). *pENTR*/*Not*I_*Asc*I is linearized *pENTR_MCS* fragment obtained through digestion with *Asc*I and *Not*I (New England Biolabs, MA, USA). *pENTR_MCS* had the multiple cloning site, including *Asc*I and *Not*I, at the cloning site of *pENTR-D-TOPO*.

A set of the three DNA fragments was integrated into each construct by the In-fusion method (In-Fusion® Snap Assembly Master Mix; Takara Bio). The *CaMV35S* promoter sequence was then cloned into the *pENTR5’-TOPO* cloning vector (Thermo Fisher Scientific). Finally, *pENTR_LUCX_AA_Ins24bp/26bp* (*att*L1-*att*L2) fragments were individually integrated with *pENTR5’_CaMV35S* (*att*L4-*att*R1) into a *pUC18*-based destination vector (*att*R4-*att*R2), carrying a *Nos* terminator, through LR reactions (Gateway LR Clonase II Enzyme mix; Thermo Fisher Scientific) [28]. Similarly, the *AtCCA1* promoter sequence was cloned into the *pENTR5’-TOPO* cloning vector [4], and *pENTR_LUC40Ins24bp/26bp* (*att*L1-*att*L2) fragments were individually integrated with *pENTR5’_AtCCA1* (*att*L4-*att*R1) into a *pUC18*-based destination vector (*att*R4-*att*R2), carrying a *Nos* terminator, through LR reactions.

*pENTR_AtU6-26::sgRNA-LUCX_AA_* was constructed as follows: The DNA fragment comprising the 20-bp target region was made by annealing fragments 5’-attgGTTCTACCTATGATTCCCAG-3’ and 5’-aaacCTGGGAATCATAGGTAGAAC-3’. This DNA fragment was cloned into the *Bbs*I-site of the *pENTR_AtU6-26::sgRNA* vector [30]. The mixture of *pENTR_AtU6-26::sgRNA-LUCX_AA_* and the plasmid expressing Cas9 (*pUC18_PcUBQ4-2::Cas9*) [47] were defined as the *CRISPR/Cas9* constructs in this study.

*pR4GWB501_CaMV35S::LUC40Ins26bp* was constructed by integrating *pENTR_LUC40Ins26bp*, *pENTR5’_CaMV35S*, and *pR4GWB501* by LR reaction. *pR4GWB501* is a binary transformation vector harboring a hygromycin-resistant selectable marker [59].

### Production of transgenic lines

Stable transformation of *Arabidopsis thaliana* (*Col-0*) with *Agrobacterium tumefaciens* (EHA105 strain) carrying *pR4GWB501_CaMV35S::LUC40Ins26bp* was performed using the floral dip method [60]. T1 seeds were sown on 0.5× MS agar medium containing 20 mg/mL hygromycin B (FUJIFILM Wako Pure Chemical Corporation, Osaka, Japan) and 200 mg/mL cefotaxime sodium (Claforan®; Sanofi, Paris, France). The 41 lines of seedlings that survived in the selection medium were transplanted into soil and grown to obtain T2 seeds. The T2 seeds from each plant were sown on the same selection agar medium. Based on the segregation of hygromycin B resistance, 13 lines were identified as single-locus transformants. Among these lines, we selected those T2 generation lines that showed high expression levels of *LUC40Ins26bp*. Homozygous lines (T2 generation) were selected based on the segregation of hygromycin B resistance in the T3 generation. The T3 seedlings of these homozygous lines were used for bioluminescence experiments.

### Particle bombardment experiments

Reporter constructs or *CRISPR/Cas9* constructs were introduced into duckweed or *Arabidopsis* using a particle bombardment system (PDS-1000/He; Bio-Rad, Hercules, CA, USA) as previously described [30, 48]. Gold particles were prewashed and stocked in 50 % glycerol (duckweed, 1-µm diameter, 60 mg/mL; *Arabidopsis*, 0.6-µm diameter, 13 mg/mL). The number of gold particles per unit volume was equalized. The concentrations and amounts of constructs coated on the gold particles in each experiment are shown in Supplementary Table S3. After gene transfection, plants were immediately transferred to 60-mm petri dishes (IWAKI, Tokyo, Japan) containing 8 mL NF medium containing 8 µL of 0.1 mM D-luciferin.

### Bioluminescence monitoring and quantification

Bioluminescence monitoring at the single-cell level was performed as previously described with minor modifications [22, 28]. A lens (XENON 0.95/22MM C-mount; Schneider Optics, Rueil-Malmaison, France) attached to a short-pass filter (SV630; Asahi Spectra, Tokyo, Japan) to reduce delayed autofluorescence from the chloroplasts was used together with a bioluminescence imaging system consisting of an EM-CCD camera (ImagEM X2 C9100-23B; Hamamatsu Photonics, Shizuoka, Japan). Extension rings of 5 and 16.5 mm thick were used for whole-dish and close-up imaging, respectively. HC-image and HOKAWO software (Hamamatsu Photonics) were used to control the imaging system. Bioluminescence images (16-bit TIFF format) were captured twice at several exposure times with the ImagEM camera (cooled at −80 °C) at an EM gain of 1,200 after at least a 4-min dark treatment for autofluorescence decay. Image analysis and post-processing were performed using ImageJ software (LOCI, University of Wisconsin, USA). To remove the cosmic-ray spike, the minimum value for each pixel between two sequential images was used for analysis. The signal intensity of the bioluminescent spot was quantified as an integrated density of the region of interest (ROI; 6 × 6 pixels). The ROIs that could be determined from a single cell were manually selected for each image. To quantify the background noise, we selected ten ROIs (10 × 10 pixels) in the plant area without bioluminescent spots and calculated the median pixel signal intensity of each ROI. We then defined the mean value of the medians of the ten ROIs as the background value for each pixel. The bioluminescence intensities were calculated using the following formula [27]:

Number of photons per min = (signal intensity − background value) × Conversion factor / (Analog gain × EM gain × Conversion efficiency) × 60 (sec) / Exposure time (sec) = (signal intensity – background value) × 5.8 / (1 × 1200 × 0.9) × 60 (sec) / Exposure time (sec).

### RNA isolation and qPCR analysis

Total RNA was isolated from five 14-day-old *Arabidopsis* seedlings, grown under constant light conditions, using a NucleoSpin® RNA kit (Takara Bio). cDNA was synthesized using ReverTra Ace (TOYOBO). qPCR was performed using a StepOnePlus Real-Time PCR System (Thermo Fisher Scientific) with the THUNDERBIRD SYBR qPCR Mix (TOYOBO). cDNA levels were normalized to those of the housekeeping control gene, *ISOPENTENYL PYROPHOSPHATE:DIMETHYLALLYL PYROPHOSPHATE ISOMERASE 2* (*AtIPP2*). The following primers were used: 5’-TCAGGTGGCTCCCGCTGAATTG-3’ and 5’-CCGTCATCGTCTTTCCGTGCTC-3’ for *LUC40Ins26bp*, and 5’-GTATGAGTTGCTTCTCCAGCAAAG-3’ and 5’-GAGGATGGCTGCAACAAGTGT-3’ for *AtIPP2*.

## Acknowledgements

This work was supported in part by the Japan Society for the Promotion of Science KAKENHI [Grant numbers JP20K06342 (S.I.), 24H02121 (S.I.), 17KT0022 (T.O.), and JP19H03245 (T.O.)], the Japan Science and Technology Agency (JST), JST ALCA (JPMJAL1108, T.O.), and JST SATREPS (JPMJSA2004, S.I., and T.O.), and JST SPRING (JPMJSP2110, R.U.). We thank Dr. H. Puchta (Karlsruhe Institute of Technology, Karlsruhe, Germany) for providing *pDe-Cas9* and *pEn-Chimera* vectors, and Dr. M. Morikawa (University of Hokkaido, Hokkaido, Japan) for providing *L. japonica* 5512. We also thank Dr. S. Nakamura (University of Tokyo, Tokyo, Japan) for his support with the construction of *LU_UC*, and S. Honda and K. Naito for their support with the construction of *nLUC40*/*cLUC40* and bioluminescence imaging. We would like to thank Editage (www.editage.jp) for English language editing.

## Author contributions

T.O., S.I., and R.U. designed the study. R.U. performed the research and analyzed the data. S.I. and R.U. cloned plasmid constructions. T.O., S.I., and R.U. wrote the manuscript.

## Data availability statement

Cellular bioluminescence data will be deposited in KURENAI (https://repository.kulib.kyoto-u.ac.jp/dspace/index-en.html?locale=en).

## References

1. Elowitz, M. B., Levine, A. J., Siggia, E. D. & Swain, P. S. Stochastic gene expression in a single cell. Science 297, 1183–1186 (2002).

2. Eldar, A. & Elowitz, M. B. Functional roles for noise in genetic circuits. Nature 467, 167–173 (2010).

3. Kim, J. & Somers, D. Rapid assessment of gene function in the circadian clock using artificial MicroRNA in Arabidopsis mesophyll protoplasts. Plant physiol. 154, 611–21 (2010).

4. Nakamichi, N. et al. Characterization of plant circadian rhythms by employing Arabidopsis cultured cells with bioluminescence reporters. Plant Cell Physiol. 45, 57–67 (2004).

5. Pokhilko, A., et al. The clock gene circuit in *Arabidopsis* includes a repressilator with additional feedback loops. Mol. Syst. Biol. 8, 574 (2012).

6. Nohales, M. A. & Kay, S. A. Molecular mechanisms at the core of the plant circadian oscillator. Nat. Struct. Mol. Biol. 23, 1061–1069 (2016).

7. Sanchez, S. E., Rugnone, M. L. & Kay, S. A. Light perception: A matter of time. Mol. Plant 13, 363–385 (2020).

8. Yakir, E. et al. Cell autonomous and cell-type specific circadian rhythms in Arabidopsis. Plant J. 68, 520–531 (2011).

9. Takahashi, N., Hirata, Y., Aihara, K. & Mas, P. A hierarchical multi-oscillator network orchestrates the *Arabidopsis* circadian system. Cell 163, 148–159 (2015).

10. Gould, P. D. et al. Coordination of robust single cell rhythms in the *Arabidopsis* circadian clock via spatial waves of gene expression. eLife 7, e31700 (2018).

11. Dixit, R. & Cyr, R. Cell damage and reactive oxygen species production induced by fluorescence microscopy: effect on mitosis and guidelines for non-invasive fluorescence microscopy. Plant J. 36, 280–290 (2003).

12. Ettinger, A. & Wittmann, T. Fluorescence live cell imaging. Methods Cell Biol. 123, 77–94 (2014).

13. Millar, A. J., Short, S. R., Chua, N. H. & Kay, S. A. A novel circadian phenotype based on firefly luciferase expression in transgenic plants. Plant Cell 4, 1075–1087 (1992).

14. Millar, A. J., Carré, I. A., Strayer, C. A., Chua, N.-H. & Kay, S. A. Circadian clock mutants in *Arabidopsis* identified by luciferase imaging. Science 267, 1161–1163 (1995).

15. Conti, E., Franks, N. P. & Brick, P. Crystal structure of firefly luciferase throws light on a superfamily of adenylate-forming enzymes. Structure 4, 287–298 (1996).

16. Thain, S. C., Murtas, G., Lynn, J. R., McGrath, Robert. B. & Millar, A. J. The Circadian clock that controls gene expression in *Arabidopsis* is tissue specific. Plant Physiol. 130, 102–110 (2002).

17. Fukuda, H., Nakamichi, N., Hisatsune, M., Murase, H. & Mizuno, T. Synchronization of plant circadian oscillators with a phase delay effect of the vein network. Phys. Rev. Lett. 99, 098102; 10.1103/PhysRevLett.99.098102 (2007).

18. Wenden, B., Toner, D. L. K., Hodge, S. K., Grima, R. & Millar, A. J. Spontaneous spatiotemporal waves of gene expression from biological clocks in the leaf. Proc. Natl. Acad. Sci. USA 109, 6757–6762 (2012).

19. Fukuda, H., Ukai, K. & Oyama, T. Self-arrangement of cellular circadian rhythms through phase-resetting in plant roots. Phys. Rev. E 86, 041917; 10.1103/PhysRevE.86.041917 (2012).

20. Ueno, K., Ito, S. & Oyama, T. An endogenous basis for synchronisation characteristics of the circadian rhythm in proliferating Lemna minor plants. New Phytol. 233, 2203–2215 (2022).

21. Nakamura, S. & Oyama, T. Adaptive diversification in the cellular circadian behavior of *Arabidopsis* leaf- and root-derived cells. Plant Cell Physiol. 63, 421–432 (2022).

22. Muranaka, T. & Oyama, T. Heterogeneity of cellular circadian clocks in intact plants and its correction under light-dark cycles. Sci. Adv. 2, e1600500 (2016).

23. Okada, M., Muranaka, T., Ito, S. & Oyama, T. Synchrony of plant cellular circadian clocks with heterogeneous properties under light/dark cycles. Sci. Rep. 7, 317; 10.1038/s41598-017-00454-8 (2017).

24. Isoda, M. & Oyama, T. Use of a duckweed species, *Wolffiella hyalina*, for whole-plant observation of physiological behavior at the single-cell level. Plant Biotechnol. 35, 387–391 (2018).

25. Watanabe, E., Isoda, M., Muranaka, T., Ito, S. & Oyama, T. Detection of uncoupled circadian rhythms in individual cells of *Lemna minor* using a dual-Color bioluminescence monitoring system. Plant Cell Physiol. 62, 815–826 (2021).

26. Muranaka, T., Kubota, S. & Oyama, T. A single-cell bioluminescence imaging system for monitoring cellular gene expression in a plant body. Plant Cell Physiol. 54, 2085–2093 (2013).

27. Muranaka, T. & Oyama, T. Application of single-cell bioluminescent imaging to monitor circadian rhythms of individual plant cells. Bioluminescent Imaging: Methods and Protocols (ed. Ripp, S.) 231–242 (Springer US, New York, NY, 2020).

28. Watanabe, E. et al. A non-cell-autonomous circadian rhythm of bioluminescence reporter activities in individual duckweed cells. Plant Physiol. 193, 677–688 (2023).

29. Serikawa, M., Miwa, K., Kondo, T., & Oyama, T. Functional conservation of clock-related genes in flowering plants: Overexpression and RNA interference analyses of the circadian rhythm in the monocotyledon *Lemna gibba*. Plant Physiol. 146, 1952–1963 (2008).

30. Kanesaka, Y., Okada, M., Ito, S. & Oyama, T. Monitoring single-cell bioluminescence of Arabidopsis leaves to quantitatively evaluate the efficiency of a transiently introduced CRISPR/Cas9 system targeting the circadian clock gene *ELF3*. Plant Biotechnol. 36, 187–193 (2019).

31. Yamashita, T., Iida, A. & Morikawa, H. Evidence that more than 90% of *β-glucuronidase*-expressing cells after particle bombardment directly receive the foreign gene in their nucleus. Plant Physiol. 97, 829–831 (1991).

32. Wenden, B., Toner, D. L. K., Hodge, S. K., Grima, R. & Millar, A. J. Spontaneous spatiotemporal waves of gene expression from biological clocks in the leaf. Proc. Natl. Acad. Sci. USA 109, 6757–6762 (2012).

33. Muranaka, T. & Oyama, T. Monitoring circadian rhythms of individual cells in plants. J. Plant Res. 131, 15–21 (2018).

34. Sherf, B. A. & Wood, K. V. Firefly luciferase engineered for im-proved genetic reporting. Promega Notes Magazine 49, 14–21 (Promega Corporation, Madison, WI, 1994).

35. Branchini, B. R., Southworth, T. L., Khattak, N. F., Michelini, E. & Roda, A. Red- and green-emitting firefly luciferase mutants for bioluminescent reporter applications. Anal. Biochem. 345, 140–148 (2005).

36. Wang, Y., Akiyama, H., Terakado, K. & Nakatsu, T. Impact of site-directed mutant luciferase on quantitative green and orange/red emission intensities in firefly bioluminescence. Sci. Rep. 3, 2490; 10.1038/srep02490 (2013).

37. Nishiguchi, T. et al. Development of red-shifted mutants derived from luciferase of Brazilian click beetle *Pyrearinus termitilluminans*. J. Biomed. Opt. 20, 101205; 10.1117/1.JBO.20.10.101205 (2015).

38. Greer III, L. F. & Szalay, A. A. Imaging of light emission from the expression of luciferases in living cells and organisms: a review. Luminescence 17, 43–74 (2002).

39. Horvath, P. & Barrangou, R. CRISPR/Cas, the Immune System of Bacteria and Archaea. Science 327, 167–170 (2010).

40. Doudna, J. A. & Charpentier, E. The new frontier of genome engineering with CRISPR-Cas9. Science 346, 1258096 (2014).

41. Zhang, F., Wen, Y. & Guo, X. CRISPR/Cas9 for genome editing: progress, implications and challenges. Hum. Mol. Genet. 23, R40–R46 (2014).

42. McVey, M., LaRocque, J. R., Adams, M. D. & Sekelsky, J. J. Formation of deletions during double-strand break repair in *Drosophila* DmBlm mutants occurs after strand invasion. Proc. Natl. Acad. Sci. USA 101, 15694–15699 (2004).

43. Tuladhar, R. et al. CRISPR-Cas9-based mutagenesis frequently provokes on-target mRNA misregulation. Nat. Commun. 10, 4056; 10.1038/s41467-019-12028-5 (2019).

44. Feng, Z. et al. Multigeneration analysis reveals the inheritance, specificity, and patterns of CRISPR/Cas-induced gene modifications in Arabidopsis. Proc. Natl. Acad. Sci. USA 111, 4632–4637 (2014).

45. Bortesi, L. & Fischer, R. The CRISPR/Cas9 system for plant genome editing and beyond. Biotechnol. Adv. 33, 41–52 (2015).

46. Butler, N. M., Baltes, N. J., Voytas, D. F. & Douches, D. S. Geminivirus-mediated genome editing in Potato (*Solanum tuberosum* L.) using sequence-specific nucleases. Front. Plant Sci. 7, 1045 (2016).

47. Fauser, F., Schiml, S. & Puchta, H. Both CRISPR/Cas-based nucleases and nickases can be used efficiently for genome engineering in *Arabidopsis thaliana*. Plant J. 79, 348–359 (2014).

48. Muranaka, T., Okada, M., Yomo, J., Kubota, S. & Oyama, T. Characterisation of circadian rhythms of various duckweeds. Plant Biol. 17, 66–74 (2015).

49. Kawade, K. & Tsukaya, H. Probing the stochastic property of endoreduplication in cell size determination of *Arabidopsis thaliana* leaf epidermal tissue. PLOS ONE 12, e0185050 (2017).

50. Jiang, S. et al. The UBP14-CDKB1;1-CDKG2 cascade controls endoreduplication and cell growth in Arabidopsis. Plant Cell 34, 1308–1325 (2022).

51. Li, H. L., Gee, P., Ishida, K. & Hotta, A. Efficient genomic correction methods in human iPS cells using CRISPR–Cas9 system. Methods 101, 27–35 (2016).

52. Kato-Inui, T., Takahashi, G., Hsu, S. & Miyaoka, Y. Clustered regularly interspaced short palindromic repeats (CRISPR)/CRISPR-associated protein 9 with improved proof-reading enhances homology-directed repair. Nucleic Acids Res. 46, 4677–4688 (2018).

53. Nakatsu, T. et al. Structural basis for the spectral difference in luciferase bioluminescence. Nature 440, 372–376 (2006).

54. Lück, S., Thurley, K., Thaben, P. F. & Westermark, P. O. Rhythmic degradation explains and unifies circadian transcriptome and proteome data. Cell Rep. 9, 741–751 (2014).

55. Gout. J. -F., Thomas, W. K., Smith, Z., Okamoto, K. & Lynch, M. Large-scale detection of in vivo transcription errors. Proc. Natl. Acad. Sci. USA 110, 46 (2013).

56. Ben-Tov, D. et al. Uncovering the dynamics of precise repair at CRISPR/Cas9-induced double-strand breaks. Nat. Commun. 15, 5096 (2024).

57. Horikawa, Y., Watanabe, E., Ito, S. & Oyama, T. Model-based analysis of the circadian rhythm generation of bioluminescence reporter activity in duckweed. Plant Biotechnol. in press; 10.1101/2024.05.26.595939 (2024).

58. Lindsey III, B. E., Rivero, L., Calhoun, C. S., Grotewold, E. & Brkljacic, J. Standardized Method for High-throughput Sterilization of Arabidopsis Seeds. J. of Vis. Exp. 128, e56587; 10.3791/56587 (2017).

59. Nakagawa, T. et al., Development of R4 Gateway Binary Vectors (R4pGWB) Enabling High-Throughput Promoter Swapping for Plant Research. Biosci. Biotech. Bioch. 72, 624–629 (2008).

60. Clough, S. J. & Bent, A. F. Floral dip: a simplified method for Agrobacterium - mediated transformation of *Arabidopsis thaliana*. Plant J. 16, 735–743 (1998).

