## Supplementary information for "A CRISPR/Cas9-induced restoration of bioluminescence reporter system for single-cell gene expression analysis in plants"

Tokitaka Oyama, Ph. D.


This file includes:

Supplementary Table 1-3

Supplementary Fig. 1-10

**Supplementary Table S1. The primers used for PCR amplification to construct *pENTR_LUC2Ins24bp/26bp***

| forward primers (from 5′ to 3′) | |
|---|---|
| *LUC2Ins24bp* | CTATGATTCCCAGCGGCACCGACGCCAAAAACATAAAGAAAGGC |
| *LUC2Ins26bp* | CTATGATTCCCAGCGGTTAACCGACGCCAAAAACATAAAGAAAGGC |
| reverse primer (from 5′ to 3′) | |
| *LUC2Ins24bp/26bp* | CCGCTGGGAATCATAGGTAGAACGACCATGGCGGCCGCGGAGCCTGC |

**Supplementary Table S2. The primers used for PCR amplification to construct *pENTR_LUCX$_{AA}$Ins24bp/26bp* except for *pENTR_LUC2Ins24bp/26bp***

| reverse primers for the *5′-part-LUC* fragment (from 5′ to 3′) | |
|---|---|
| *LUC40Ins24bp/26bp* | CCGCTGGGAATCATAGGTAGAACTGTTCCAGGAACCAGGGCGT |
| *LUC106Ins24bp/26bp* | CCGCTGGGAATCATAGGTAGAACCGCGGGCGCAACTGCAACTC |
| *LUC239Ins24bp/26bp* | CCGCTGGGAATCATAGGTAGAACTAAAATCGCAGTATCCGGAA |
| *LUC378Ins24bp/26bp* | CCGCTGGGAATCATAGGTAGAACATCCAGATCCACAACCTTCG |
| *LUC491Ins24bp/26bp* | CCGCTGGGAATCATAGGTAGAACTCCGTGCTCCAAAACAACAA |
| forward primers for the *3′-part-LUC* fragment (from 5′ to 3′) | |
| *LUC40Ins24bp* | GTTCTACCTATGATTCCCAGCGGCATTGCTTTTACAGATGCACA |
| *LUC40Ins26bp* | GTTCTACCTATGATTCCCAGCGGTTAATTGCTTTTACAGATGCACA |
| *LUC106Ins24bp* | GTTCTACCTATGATTCCCAGCGGCAACGACATTTATAATGAACG |
| *LUC106Ins26bp* | GTTCTACCTATGATTCCCAGCGGTTAAACGACATTTATAATGAACG |
| *LUC239Ins24bp* | GTTCTACCTATGATTCCCAGCGGCAGTGTTGTTCCATTCCATCA |
| *LUC239Ins26bp* | GTTCTACCTATGATTCCCAGCGGTTAAGTGTTGTTCCATTCCATCA |
| *LUC378Ins24bp* | GTTCTACCTATGATTCCCAGCGGCACCGGGAAAACGCTGGGCGT |
| *LUC378Ins26bp* | GTTCTACCTATGATTCCCAGCGGTTAACCGGGAAAACGCTGGGCGT |
| *LUC491Ins24bp* | GTTCTACCTATGATTCCCAGCGGCAAGACGATGACGGAAAAAGA |
| *LUC491Ins26bp* | GTTCTACCTATGATTCCCAGCGGTTAAAGACGATGACGGAAAAAGA |

**Supplementary Table S3. Plasmid concentrations and amounts used for particle bombardment**

| plasmid components | | | concentration (ng/µL) | amount added (µL) |
|---|---|---|---|---|
| vector | promoter | gene | | |
| *pUC18* | *CaMV35S* | *LUC2Ins24bp* | 1023.5 | 2.0 |
| | | *LUC40Ins24bp* | 989.7 | 2.0 |
| | | *LUC106Ins24bp* | 998.0 | 2.0 |
| | | *LUC239Ins24bp* | 999.9 | 2.0 |
| | | *LUC378Ins24bp* | 982.0 | 2.0 |
| | | *LUC491Ins24bp* | 1018.7 | 2.0 |
| | *CaMV35S* | *LUC2Ins26bp* | 236.1 | 8.5 |
| | | *LUC40Ins26bp* | 997.4 | 2.0 |
| | | *LUC106Ins26bp* | 366.9 | 5.5 |
| | | *LUC239Ins26bp* | 355.4 | 5.6 |
| | | *LUC378Ins26bp* | 327.8 | 6.1 |
| | | *LUC491Ins26bp* | 335.8 | 6.0 |
| | *AtCCA1* | *LUC40Ins24bp* | 1008.2 | 2.0 |
| | | *LUC40Ins26bp* | 977.1 | 2.0 |
| *pENTR* | *AtU6-26* | $sgRNA\_LUCX_{AA}$ | 998.9 | 1.5 |
| | | *sgRNA* | 1000.9 | 1.5 |
| *pUC18* | *PcUBQ4-2* | *Cas9* | 1008.4 | 1.5 |

**Supplementary Fig. S1**

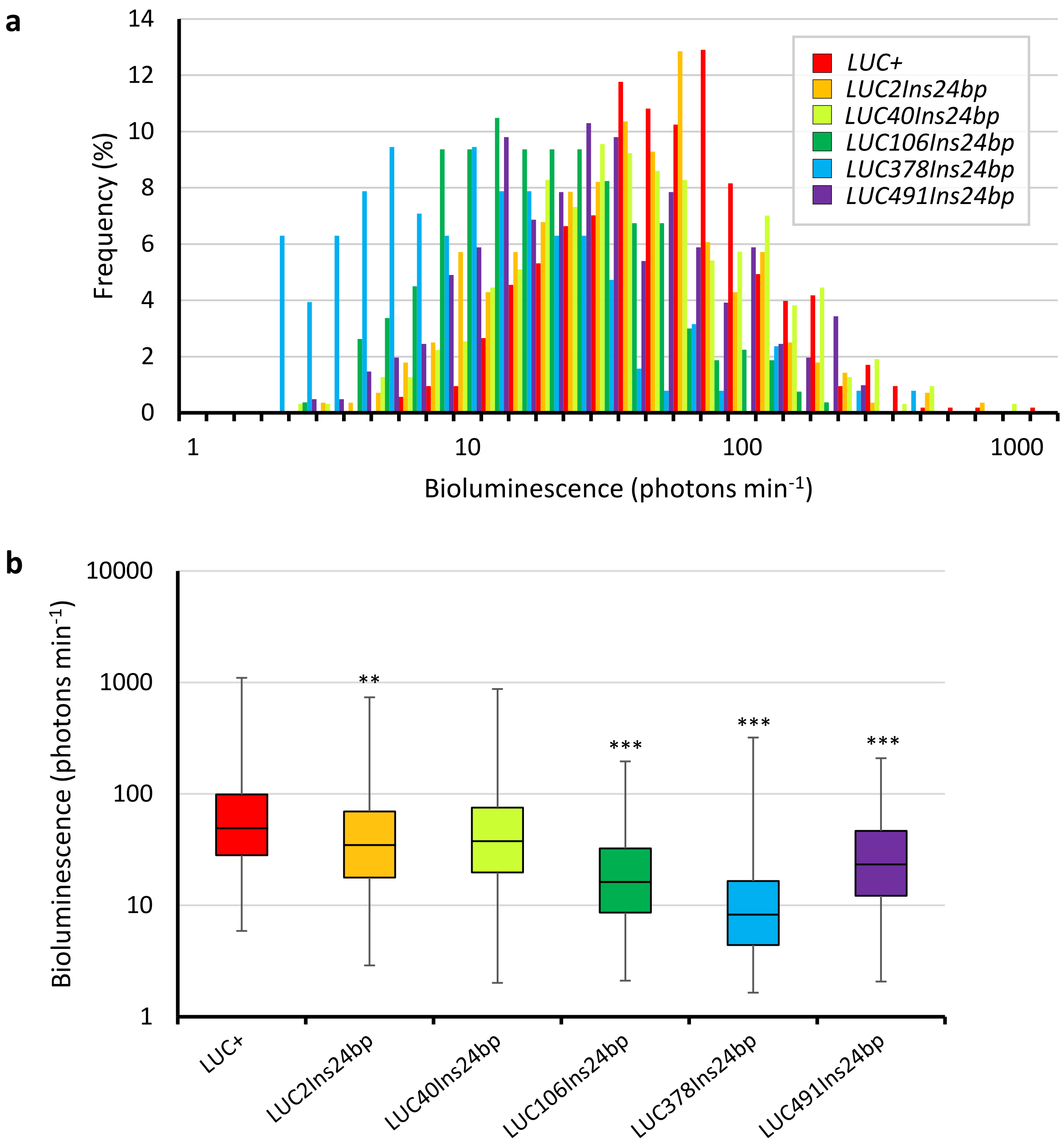


**Bioluminescence intensities of duckweed cells transfected with *LUCX$_{AA}$Ins24bp*.** (**a**) Frequency distribution (%) and (**b**) boxplot of cellular bioluminescence intensities for *CaMV35S::LUC+* (n = 526), *CaMV35S::LUC2Ins24bp* (n = 280), *CaMV35S::LUC40Ins24bp* (n = 314), *CaMV35S::LUC106Ins24bp* (n = 267), *CaMV35S::LUC378Ins24bp* (n = 127), and *CaMV35S::LUC491Ins24bp* (n = 204). Within each box for (**b**), the horizontal line represents the median value, the box extends from the 25th to the 75th percentile, and the vertical extended line ranges from the maximum to the minimum. The bioluminescence distribution for each construct is compared to that of *CaMV35S::LUC+* (Student's two-tailed test; *$p$ < 0.05, **$p$ < 0.01, ***$p$ < 0.001).

**Supplementary Fig. S2**

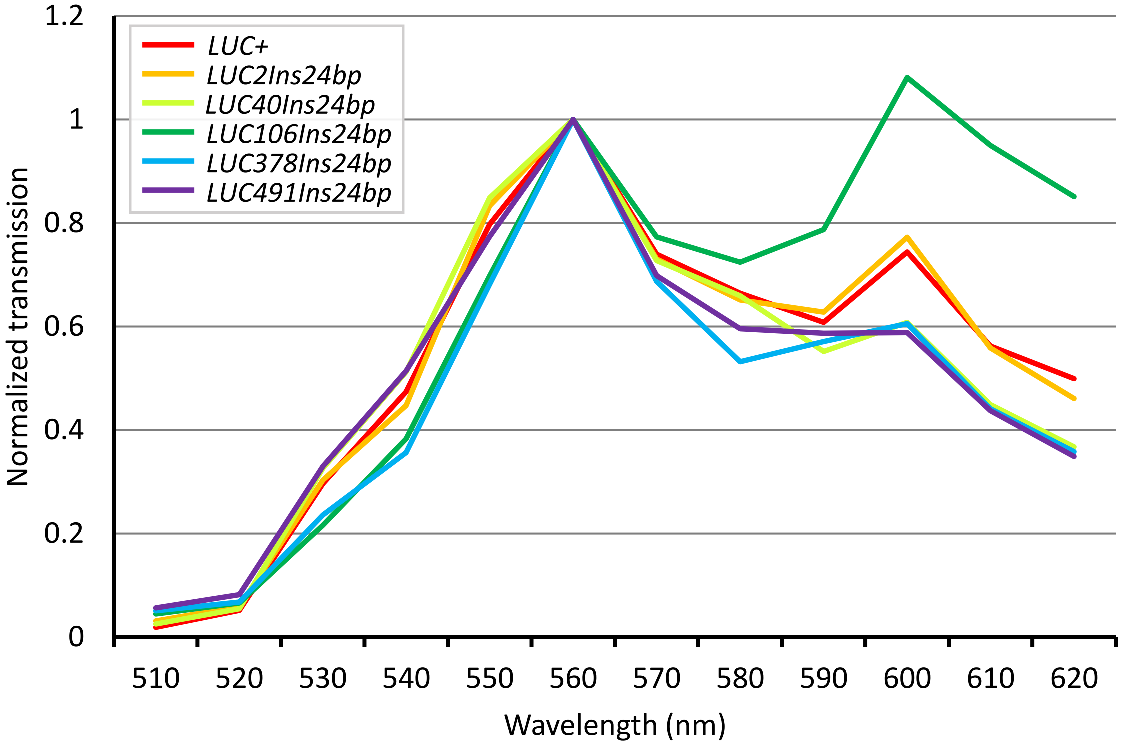


**Bioluminescence spectra of *LUCX$_{AA}$Ins24bp*.** The transmitted bioluminescence intensities from tested constructs were normalized to that at 560 nm were plotted. Bioluminescence was filtered with a series of band-pass filters and quantified.

**Supplementary Fig. S3**

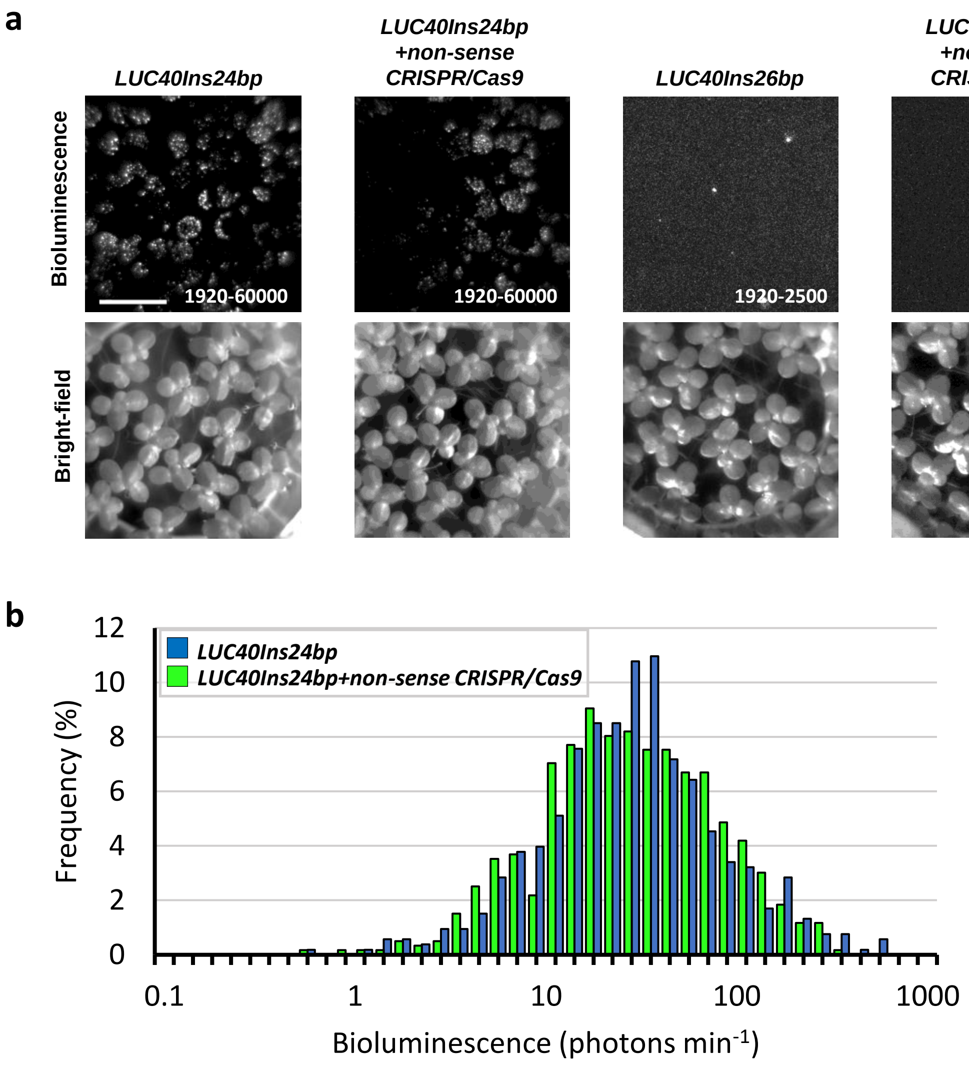


**Bioluminescence in duckweed cells co-transfected with *LUC40Ins24bp* or *LUC40Ins26bp* and nonsense-*CRISPR/Cas9* constructs.** (**a**) Bioluminescence (top) and bright-field (bottom) images of duckweed plants transfected with a reporter construct or both a reporter and nonsense-*CRISPR/Cas9* constructs (indicated above each set of images). Signal ranges are indicated in each bioluminescence image. Exposure time: 60 s. Bar: 10 mm. (**b**) Frequency distribution (%) of bioluminescence intensities in cells (co-)transfected with *CaMV35S::LUC40Ins24bp* (blue bars; n = 529) or *CaMV35S::LUC40Ins24bp* and nonsense-*CRISPR/Cas9* constructs (green bars; n = 597).

**Supplementary Fig. S4**

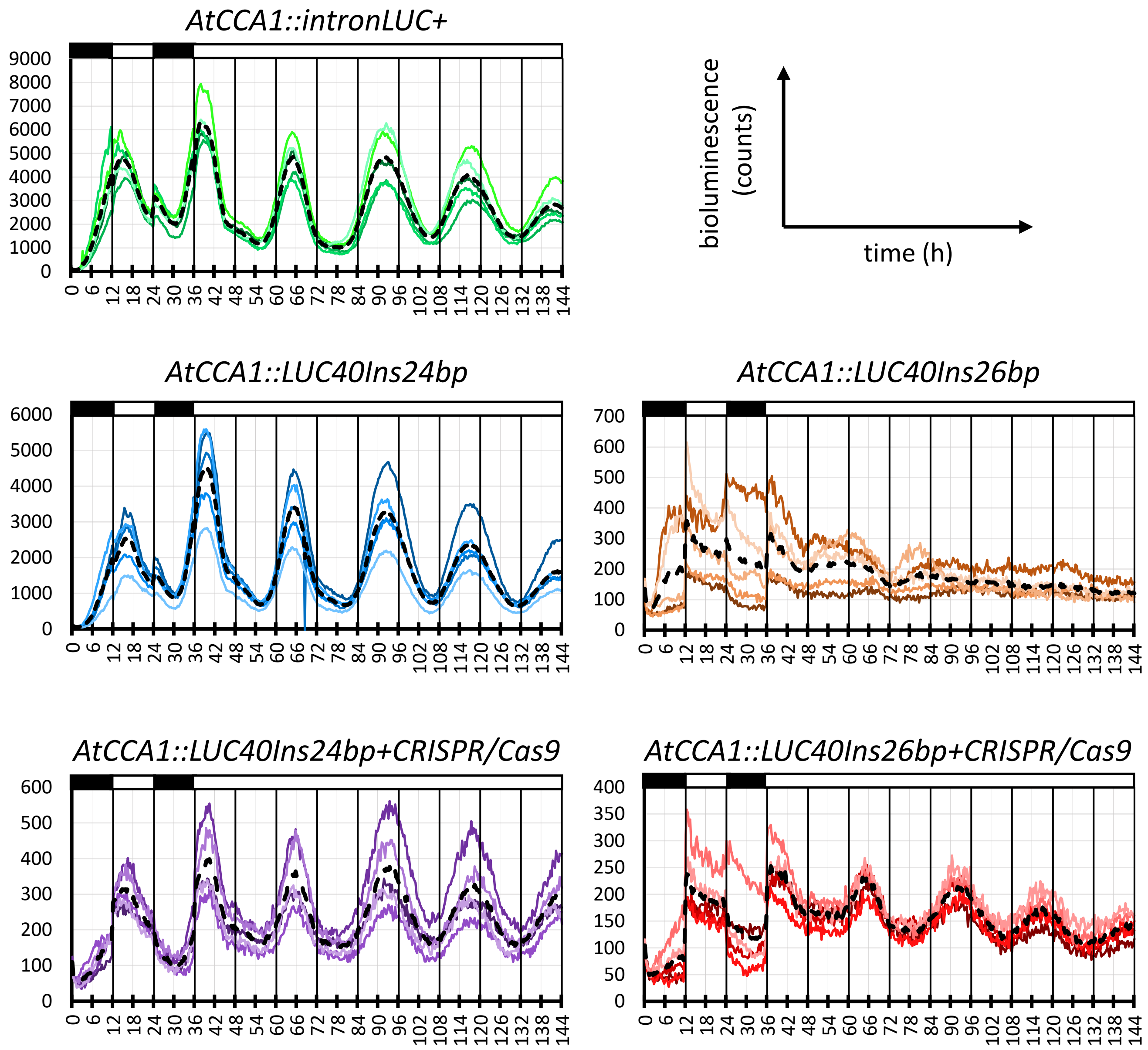


**Effects of the *CRISPR/Cas9* constructs on bioluminescence rhythms of duckweed plants transiently transfected with *AtCCA1::LUC40Ins24bp* and *AtCCA1::LUC40Ins26bp* reporters.** Each graph represents a time series of the bioluminescence intensities observed in duckweed plants transfected with a reporter construct and the *CRISPR/Cas9* construct (indicated above each graph). Black dotted lines represent the mean bioluminescence intensities of five samples. As previously described, bioluminescence was measured using an automatic luminescence monitoring system with a photomultiplier tube [48]. The *intronLUC*+ gene has an intron within the *LUC*+ coding region [48]. Black and white boxes indicate dark and light conditions, respectively.

**Supplementary Fig. S5**

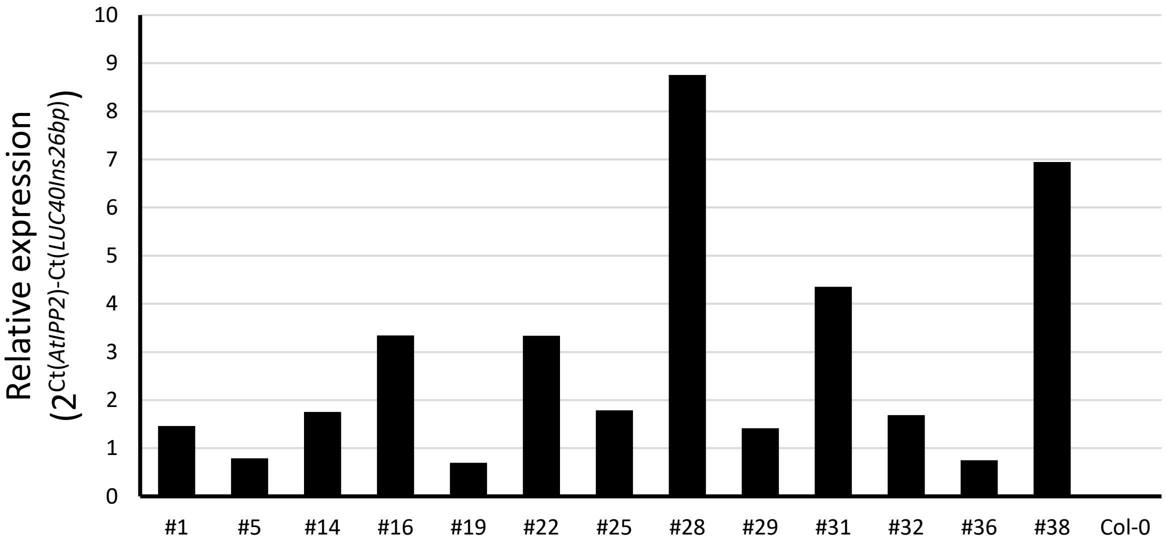


**mRNA expression levels of *LUC40Ins26bp* in T2 transgenic *Arabidopsis* plants carrying *CaMV35S::LUC40Ins26bp*.** *LUC40Ins26bp* mRNA levels, quantified by qPCR, are shown for 13 independent transgenic lines and *Col-0*. Based on the segregation of hygromycin B resistance, these lines were estimated as single-locus transformants.

### Supplementary Fig. S6

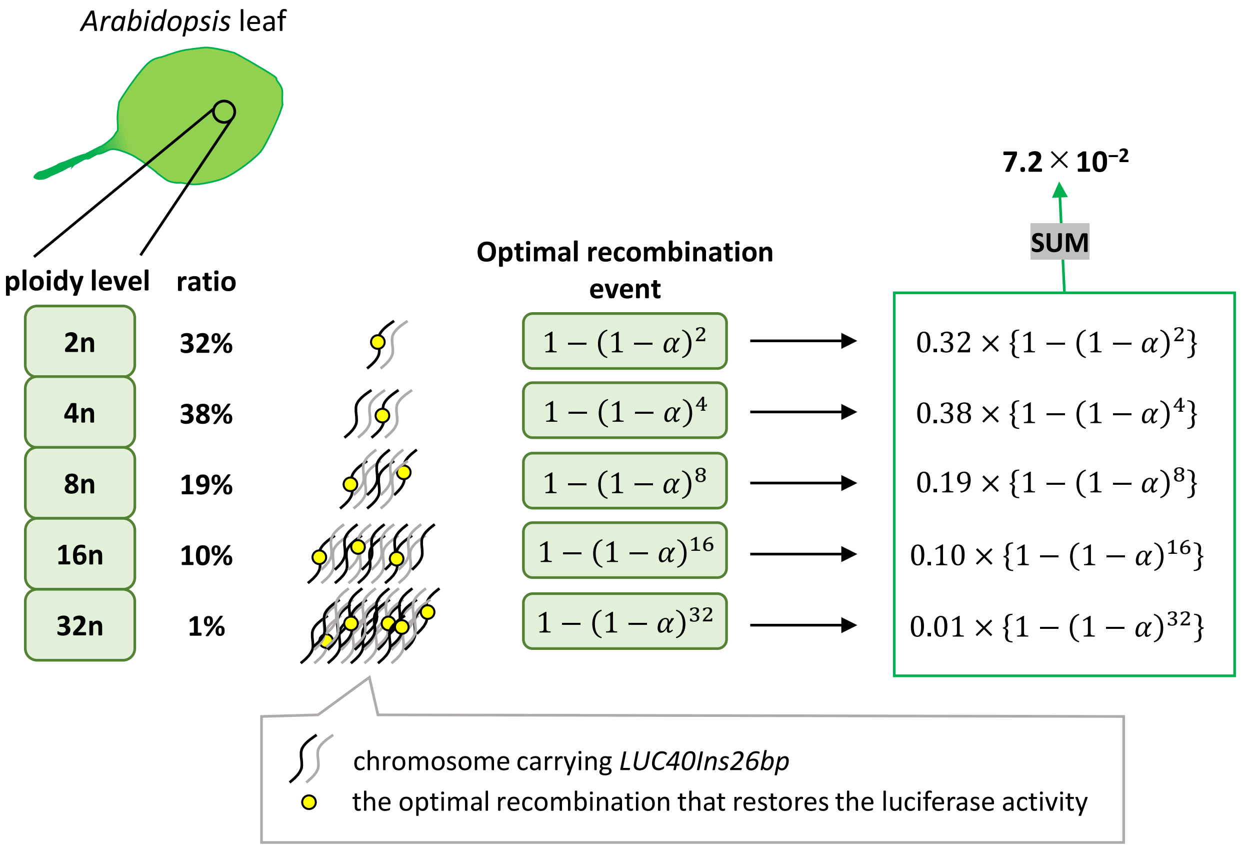


**Relationship between the ploidy profile of *Arabidopsis* leaf cells and the ratio of bioluminescent spots.** The indicated ploidy levels represent the ploidy profile of *Arabidopsis* leaf epidermal cells (2n:4n:8n:16n:32n = 0.32:0.38:0.19:0.10:0.01) reported by Kawade and Tsukaya, 2017 [49]. The probability of the optimal recombination was defined as $\alpha$. The probability of restoring cellular bioluminescence by the optimal recombination was expressed as follows;

$$1-(1-\alpha)^{(\text{ploidy level})}.$$

The ratio of bioluminescence-restored cells to transfected cells ($7.2\times10^{-2}$; Fig. 4) was simulated as the sum of the probabilities with a value of $\alpha = 1.37\times10^{-2}$.

**Supplementary Fig. S7**

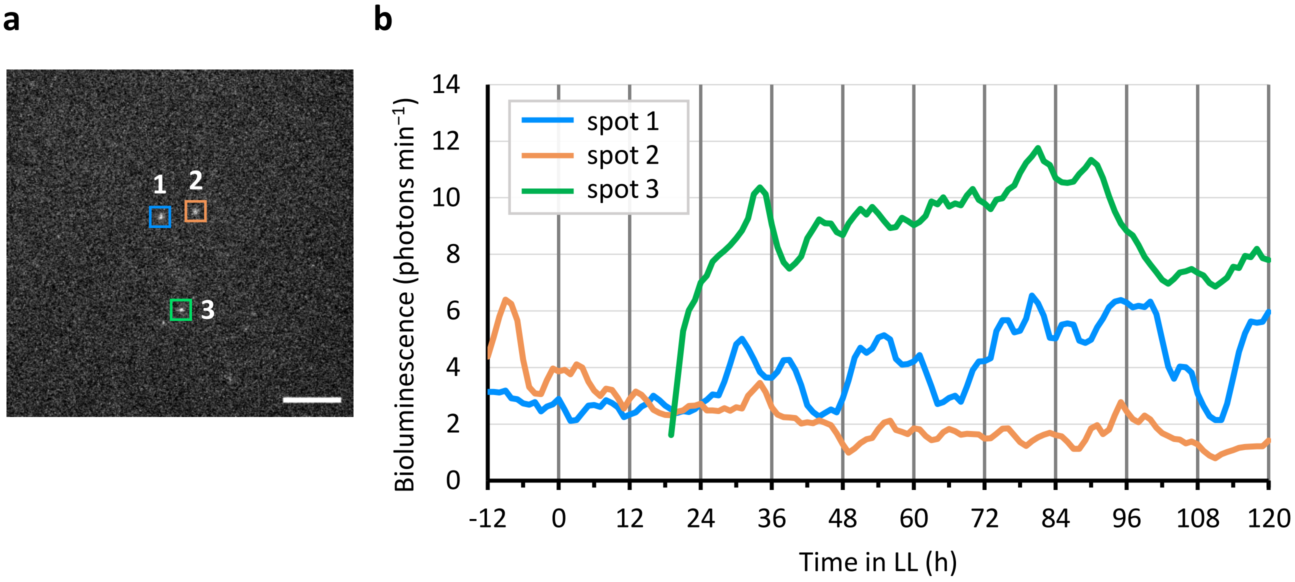


**Cellular bioluminescence traces of restored-*LUC40Ins26bp* in transgenic plants (line *LUC40Ins26bp #38-2*).** (**a**) Snapshot of bioluminescence at 28 h in constant light. Each colored square represents one bioluminescent spot of interest. Spot 3 belongs to a different leaf from that bearing spots 1 and 2. Bar: 2 mm. (**b**) Cellular bioluminescence traces of the three bioluminescent spots shown in (**a**). Quantified intensities (3-h moving average) are plotted for each spot. Time schedule was the same as Fig. 6.

### Supplementary Fig. S8

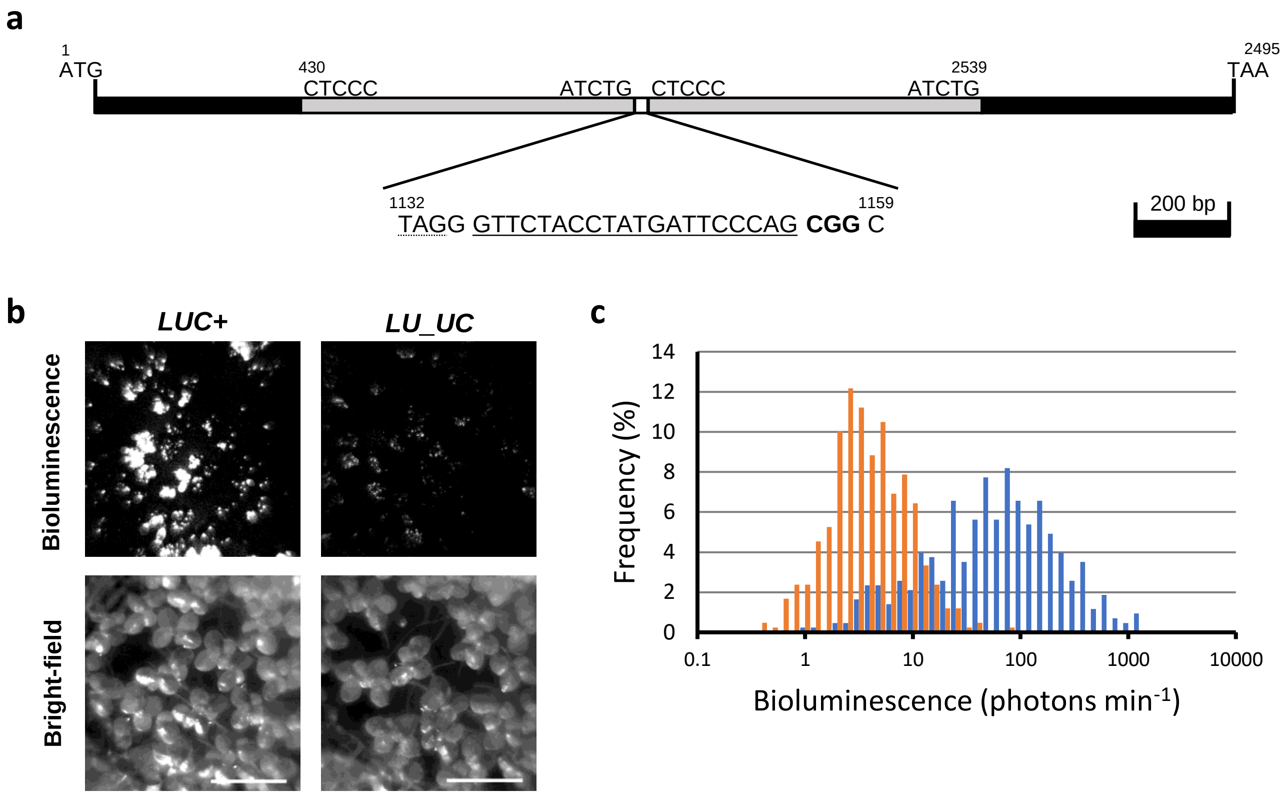


***LU_UC* construct and its bioluminescence properties.** (**a**) Map of the *LU_UC* gene. The region encompassing from 430 bp to 1131 bp of the *LUC*+ gene is duplicated. A 28-bp fragment (1132–1159 bp) was inserted between these duplicated sequences. The *sgRNA* target sequence for CRISPR/Cas9 is underlined, whereas the PAM sequence CGG is indicated in bolds. TAG, a stop codon, is underlined with a dotted line. (**b**) Bioluminescence (top) and bright-field (bottom) images of duckweed plants transfected with *CaMV35S::LUC*+ (left) and *CaMV35S::LU_UC* (right). The signal range for each bioluminescence image was fixed to 1920–5000. Exposure time: 30 s. Bars: 10 mm. (**c**) Frequency distribution (%) of cellular bioluminescence intensities for *CaMV35S::LUC*+ (blue bars, n = 427) and *CaMV35S::LU_UC* (orange bars, n = 419).

**Supplementary Fig. S9**

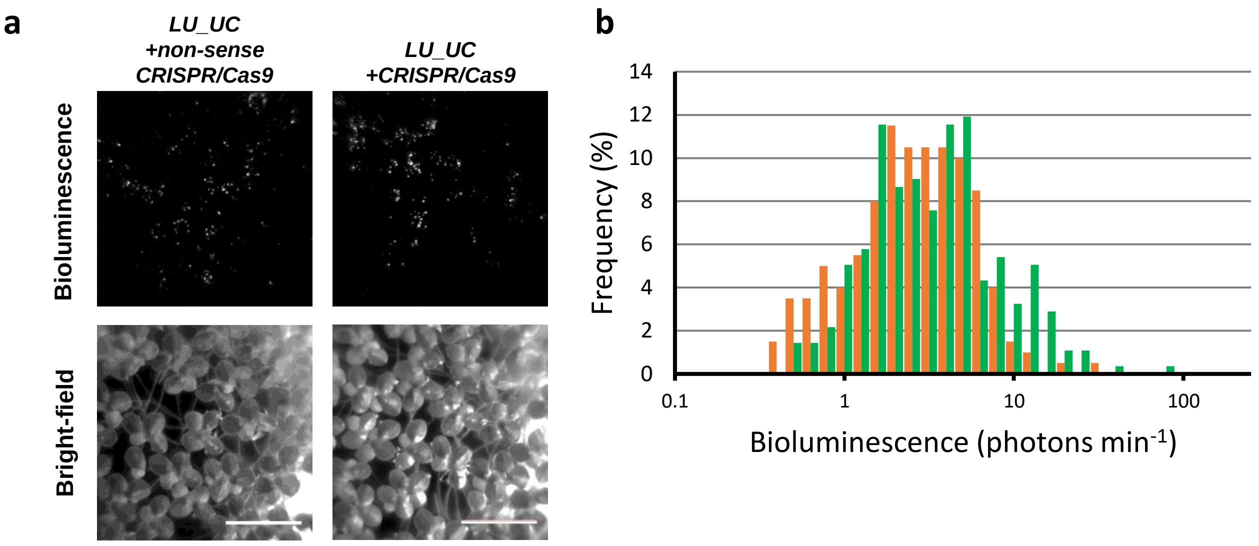


**Comparison of cellular bioluminescence intensities between duckweed cells co-transfected with *LU_UC* and nonsense-*CRISPR/Cas9* or *CRISPR/Cas9* constructs.** (**a**) Bioluminescence (top) and bright-field (bottom) images of duckweed plants co-transfected with *CaMV35S::LU_UC* and nonsense-*CRISPR/Cas9* constructs (left), and *CaMV35S::LU_UC* and *CRISPR/Cas9* constructs (right). The signal range for each bioluminescence image was fixed to 1920–5000. Exposure time: 60 s. Bars: 10 mm. (**b**) Frequency distribution (%) of cellular bioluminescence intensities for *CaMV35S::LU_UC* and nonsense-*CRISPR/Cas9* constructs (orange bars, n = 200), and *CaMV35S::LU_UC* and *CRISPR/Cas9* constructs (green bars, n = 277).

### Supplementary Fig. S10

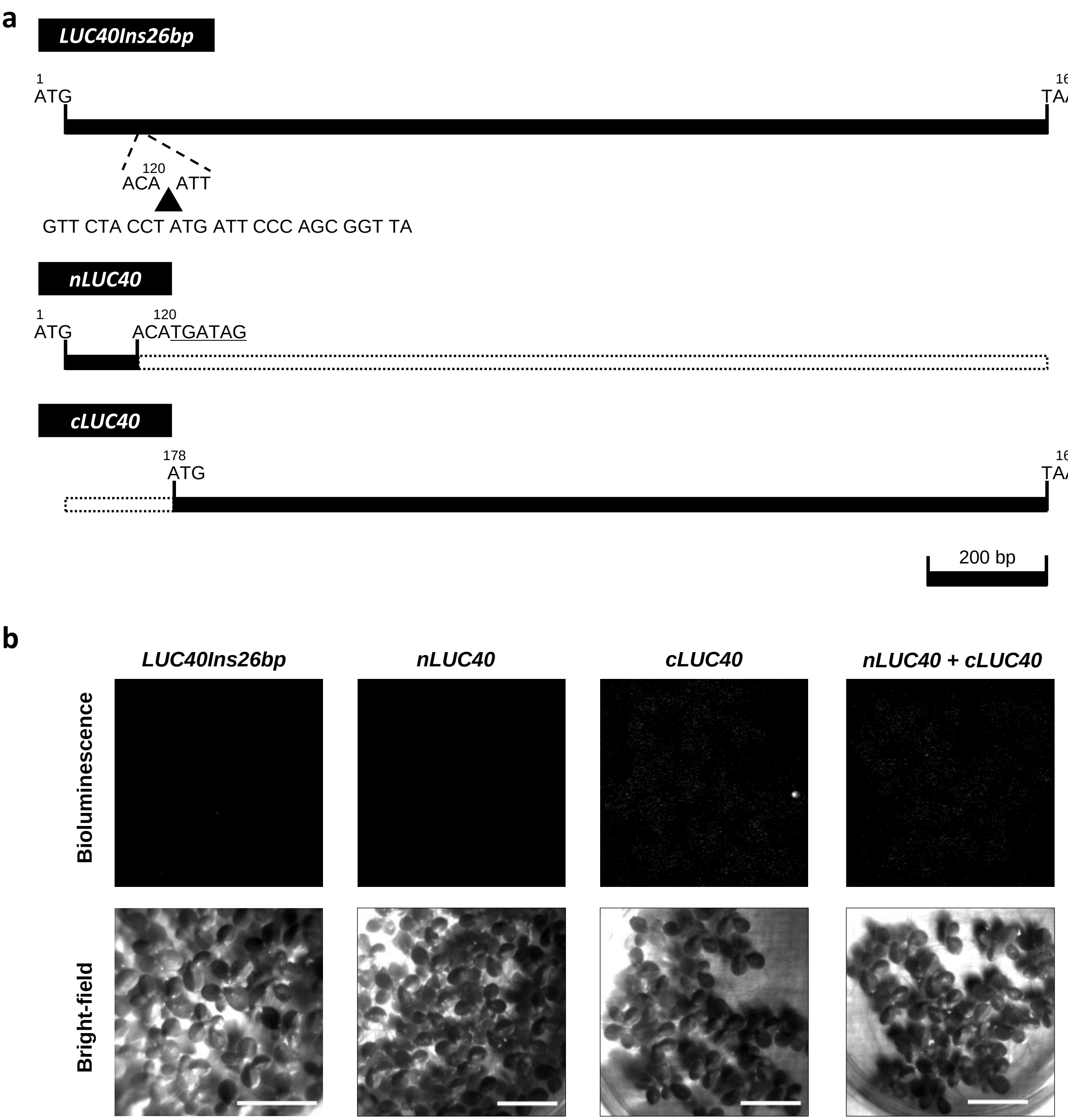


**Split *LUC40Ins24bp* constructs (*nLUC40* and *cLUC40*) and bioluminescence properties.** (**a**) *LUC40Ins26bp* (Fig. 1), *nLUC40*, and *cLUC40* gene maps. The *nLUC40* sequence has two stop codons at its end (underlined). White dotted boxes indicate deleted regions. (**b**) Bioluminescence (top) and bright-field (bottom) images of duckweed plants transfected with *CaMV35S::LUC40Ins26bp, CaMV35S::nLUC40, CaMV35S::cLUC40*, and *CaMV35S::nLUC40* + *CaMV35S::cLUC40* (from left to right). Exposure time: 90 s. The signal range for the bioluminescence images was fixed to 1920–2200. Bar: 10 mm.
